# ID1 and CEBPA Coordinate Epidermal Progenitor Cell Differentiation

**DOI:** 10.1101/2022.03.09.483653

**Authors:** Christina Geraldine Kantzer, Wei Yang, David Grommisch, Kim Vikhe Patil, Kylie Hin-Man Mak, Maria Genander

## Abstract

The regulatory circuits that coordinate epidermal differentiation during development are still not fully understood. Here we report that the transcriptional regulator ID1 is enriched in basal epidermal progenitor cells and find ID1 expression to be diminished upon differentiation. *In utero* silencing of *Id1* impairs progenitor cell proliferation, leads to precocious delamination of targeted progenitor cells and enables differentiated keratinocytes to retain progenitor markers and characteristics. Transcriptional profiling suggests ID1 acts by mediating adhesion to the basement membrane while inhibiting spinous layer differentiation. Co-immunoprecipitation reveals ID1 binding to transcriptional regulators of the class I bHLH family. We localize bHLH *Tcf3, Tcf4* and *Tcf12* to epidermal progenitor cells during epidermal stratification and established TCF3 as a downstream effector of ID1-mediated epidermal proliferation. Finally, we identify crosstalk between CEBPA, a known mediator of epidermal differentiation, and *Id1* and demonstrate that CEBPA antagonizes BMP-induced activation of *Id1.* Our work establishes ID1 as a key coordinator of epidermal development, acting to balance progenitor proliferation with differentiation and unveils how functional crosstalk between CEBPA and *Id1* orchestrates epidermal lineage progression.

## Introduction

Progenitor cells in self-renewing tissues commonly undergo a coordinated program of differentiation to accommodate tissue function. Although many aspects of progenitor cell differentiation are known, the transcriptional programs that orchestrate epidermal differentiation during development are not well defined. Epidermal p63^+^ progenitor cells are specified from a single-layered K8/K18^+^ ectoderm at embryonic day 8.5 (E8.5) (Zhao et al., 2015). Following this initial epidermal specification, progenitors commit to stratification and a transient proliferative yet suprabasal cell population co-expressing K5/K14 progenitor and K10 differentiation marker forms (Koster and Roop, 2007). At around E15, this intermediate cell population initiate terminal differentiation and proliferation becomes restricted to the basal progenitor layer (Blanpain and Fuchs, 2006). Continued growth of the embryo and expansion of the stratified embryonic epidermis thus requires coordination of progenitor cell cycle exit, delamination and differentiation through transcriptional effectors acting to either promote (Blanpain et al., 2006; Klein et al., 2017; Okuyama et al., 2004) or repress (Bao et al., 2013; Mulder et al., 2012) epidermal differentiation. It is likely that seemingly antagonistic transcriptional programs are interdependent (Boxer et al., 2014; Hopkin et al., 2012) and enable fine-tuning of epidermal lineage progression.

Several families of secreted signaling molecules have been implicated in skin development. Epidermal BMP (bone morphogenic protein) signaling is complex (Blessing et al., 1996), but collective evidence suggests a role in inhibiting progenitor cell proliferation and promoting differentiation through activation of pSMAD1/5 transcriptional programs (Botchkarev and Sharov, 2004). The ID (Inhibitors of Differentiation) proteins are established BMP-sensing target genes in several cellular contexts (Korchynskyi and ten Dijke, 2002; Kowanetz et al., 2004), including the hair follicle stem cell lineage (Genander et al., 2014). ID proteins are typically expressed in stem and progenitor cells only to be down regulated upon differentiation (Lasorella et al., 2001), suggesting - paradoxically from an epidermal BMP signaling perspective - that ID proteins could, as the name implies, enable progenitor selfrenewal by suppressing differentiation.

There are four ID family members (ID1 to ID4) all belonging to the basic helix-loop-helix (bHLH) group of transcription factors (Benezra et al., 1990). ID proteins display high bHLH domain sequence homology, rendering them capable of binding and dimerizing with other bHLH proteins. ID proteins however lack a DNA binding domain, thereby acting to inhibit functional dimerization and transcriptional activity of interacting bHLH proteins (Benezra et al., 1990; Norton, 2000). Although the most well characterized ID binding partners are E-box binding class I bHLH transcription factors such as TCF3 (E2A), TCF4 (E2.2) and TCF12 (HEB), ID proteins also exert inhibitory functions towards other transcription factor families and interact directly with cell cycle regulators (Iavarone et al., 1994; Roberts et al., 2001; Yates et al., 1999).

Expression of *Id1* is associated with stemness in several cellular contexts (Jankovic et al., 2007; Liu et al., 2013; Zhang et al., 2014). Embryonic stem cell pluripotency is sustained by BMP4-mediated expression of ID1 (Ying et al., 2003) and loss of ID1 affects both pluripotency and differentiation factors (Malaguti et al., 2013; Romero-Lanman et al., 2012). Neural stem cells require *Id1/Id3* to maintain self-renewal capacity and anchorage of stem cells to the niche (Nam and Benezra, 2009; Niola et al., 2012) through bHLH target genes. In contrast, ID1 mediate hair follicle stem cell quiescence and is required for hair follicle progenitor cell specification (Genander et al., 2014), indicating that the phenotypic outcome of ID1 is context dependent and likely a reflection of available interacting partner(s). Forced expression of ID1 in immortalized human HaCaT organotypic cultures result in a hyperproliferative, disorganized epidermis (Rotzer et al., 2006) and ID1 is highly expressed in human psoriatic skin (Bjorntorp et al., 2003) suggesting that increased ID1 levels disturb the balance between adult epidermal progenitor cell proliferation and differentiation.

The CEBPA and CEBPB transcription factors (CCAAT/enhancer binding proteins) are induced upon epidermal differentiation, and couple cell cycle exit with commitment to differentiation in several cell types, including the epidermis (Loomis et al., 2007; Lopez et al., 2009; Oh and Smart, 1998; Zhu et al., 1999). In the hematopoietic lineage, CEBPs are able to redirect chromatin binding of general signalling pathway mediators, including SMAD proteins, thereby acting to define cellular identity during lineage specification and differentiation (Trompouki et al., 2011). In other systems, SMADs bind and inhibit the transcriptional activity of CEBPs (Coyle-Rink et al., 2002; Zauberman et al., 2001), suggesting that crosstalk between SMADs and CEBPs is a commonly employed mechanism in place to fine-tune gene regulation.

Here we explored published single-cell profiling datasets (Fan et al., 2018) to identify enrichment of *Id1* in E13 progenitor cells committed to differentiation. Employing an *in utero* lentiviral injection strategy (Beronja et al., 2010) to target *in vivo* epidermal progenitors in combination with transcriptional profiling of cultured epidermal progenitor cells allowed us to delineate the consequences of *Id1* silencing on progenitor proliferation and differentiation during the onset of epidermal stratification. We demonstrate that targeting of ID1 leads to a thinning of the developing epidermis, impairs epidermal progenitor proliferation and results in loss of *Id1* silenced progenitor cells. Furthermore, *in vivo* targeted cells co-express progenitor (K5) and differentiation (K10) markers and transcriptional profiling reveals upregulation of differentiation markers *in vitro*, indicating that ID1 acts to repress epidermal differentiation in progenitor cells.

Co-immunoprecipitation demonstrates that ID1 binds the bHLH TCF family of transcription factors, and we visualize the presence of *Tcf3, Tcf4* and *Tcf12* in epidermal progenitor and committed keratinocytes during epidermal stratification. We continue to establish TCF3 as a repressor of epidermal progenitor cell proliferation *in vitro.* Focusing on CEBPA as an ID1-dependent regulator of epidermal cell cycle exit and differentiation *in vivo,* we confirm upregulation of CEBPA after silencing of *Id1* in the epidermis but fail to conclusively establish CEBPA as a bHLH target gene. Finally, we unearth a new role for CEBPA in antagonizing BMP-induced *Id1* promoter activation, establishing a regulatory mechanism of ID1 expression in epidermal progenitor cells likely to be relevant for other BMP-sensing genes. Collectively, these data establish ID1 as an essential orchestrator of epidermal differentiation and demonstrate functional crosstalk between CEBPA and ID1, enabling coordinated epidermal progenitor cell differentiation.

## Results

### ID1 is expressed in epidermal progenitor cells during skin development

Although epidermal development has been studied in some detail (Flora and Ezhkova, 2020; Miroshnikova et al., 2019; Miyai et al., 2016; Soares and Zhou, 2018), the process of epidermal differentiation is not fully understood. We exploited a published single-cell RNA-sequencing data set (Fan et al., 2018) from E13 mouse epidermis, a developmental time point where epidermal progenitor cells have not yet initiated terminal differentiation. Re-analysis revealed two main epidermal clusters, where cluster 1 was marked by the progenitor cell marker *Krt15,* and differentiation-associated *Krtdap* segregated to cluster 2 (Figure S1A-S1C). Gene Ontology (GO) profiling of differentially expressed genes (DEGs) showed enrichment of biological processes linked to tissue and epithelial development in cluster 1, whereas cluster 2 DEGs were associated with epithelial differentiation (Figure S1D) suggesting that cluster 2 epidermal cells have committed to epidermal differentiation. Interestingly, classification of protein types found in the most significant GO terms in cluster 1 and 2 revealed that whereas tissue development correlates to metabolic enzymes and signaling molecules, epithelial differentiation is associated with DNA-binding transcriptional regulators and cytoskeletal rearrangement (Figure S1E). Collectively, these results highlight the importance of transcriptional regulation in epidermal commitment to differentiation.

In addition to know transcriptional regulators such as *Mafb, Hes1* and *Klf4* (Blanpain et al., 2006; Lopez-Pajares et al., 2015; Segre et al., 1999), we identified *Id1* to be enriched in cluster 2, although prominently expressed also in cluster 1 (Figure S1F and 1A-B). Localization of ID1 protein in the developing epidermis revealed ID immunoreactivity in epidermal progenitors at E14.5 in line with the single-cell RNA-sequencing expression data from E13 (Figure 1C). Starting from E15.5, epidermal ID1 expression becomes enriched in basal progenitor cells and immunoreactivity is gradually diminished as progenitors delaminate from the basal layer and commit to differentiation (Figure 1C and S1G). Analogous to the ID1 *in vivo* expression pattern, cultured primary epidermal progenitors down regulate *Id1* mRNA and corresponding protein when asked to differentiate *in vitro* (Figure 1D and 1E). Interestingly, ID1 protein levels are sustained during the first 24 hours of differentiation, indicating a role of ID1 in the transition from basal epidermal progenitor to committed epidermal keratinocyte.

**Figure 1:**
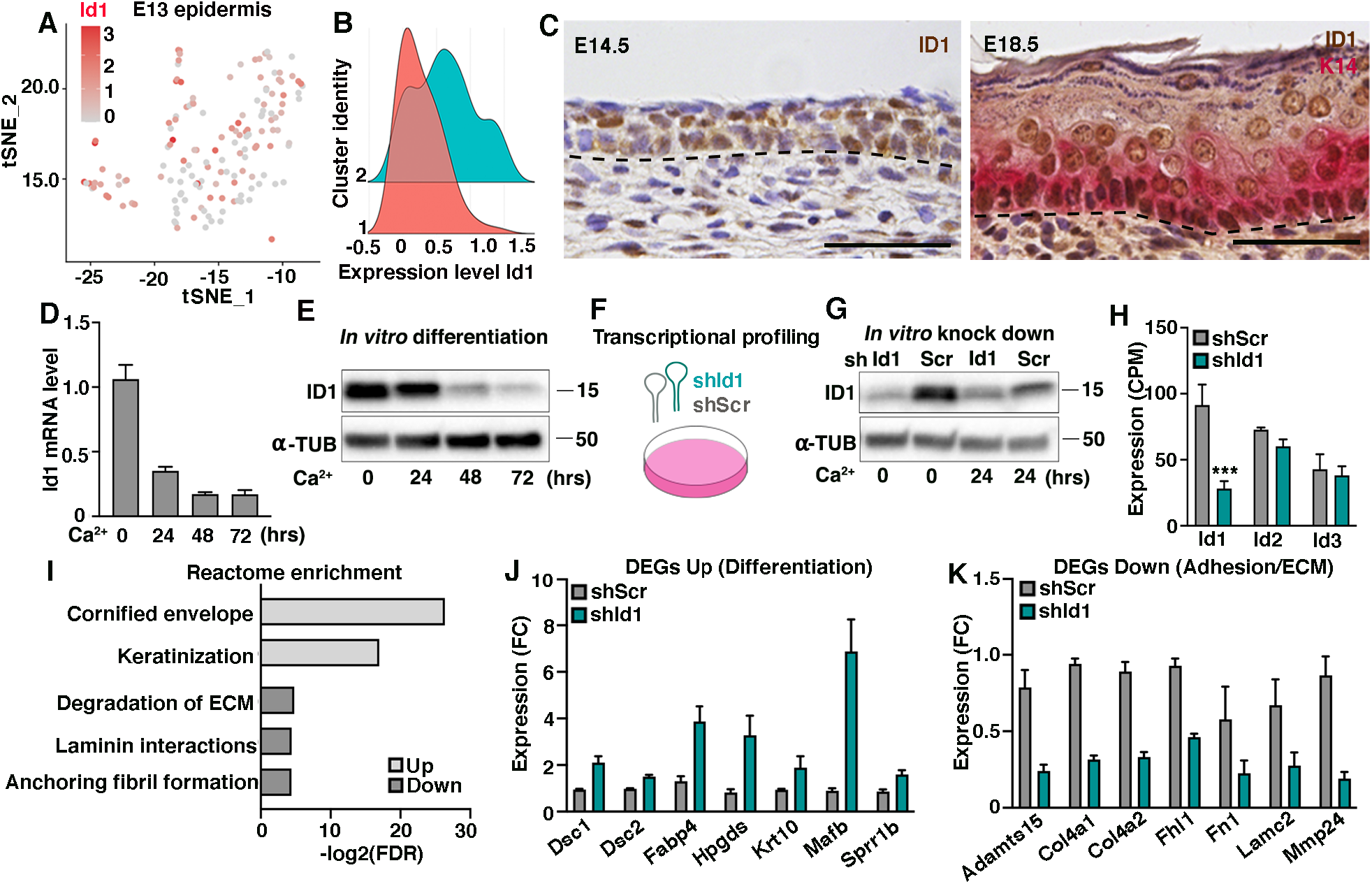
ID1 is expressed in epidermal progenitor cells during skin development. (A) Feature plot displaying expression of *Id1* in E13 epidermis analyzed by single-cell RNA-sequencing. (B) *Id1* expression in cluster 1 and cluster 2 at E13. (C) ID1 is abundantly expressed in the E14.5 developing epidermis. At E18.5, ID1 immunoreactivity is enriched in basal progenitors with decreasing expression in suprabasal layers. (D) *Id1* mRNA expression decrease with differentiation of primary epidermal progenitor cells *in vitro*. (E) ID1 protein expression dynamics during *in vitro* differentiation. (F) Schematics illustrating the transcriptional profiling exploiting shRNAs targeting scrambled control (*shScr)* or Id1 (*shId1*) mRNA sequences. (G) Western blot visualizing ID1 protein levels after *shId1* or *shScr* targeting of epidermal progenitors at 0 and 24 hours of differentiation. (H) Expression of *Id2* and *Id3* is not altered upon *shId1* targeting compared to *shScr* in epidermal progenitors displayed as CPM (counts per million) from RNA-sequencing data. (I) Reactome enrichment analysis of genes differentially expressed (>2xFC) in *shId1* compared to *shScr* progenitors. (J and K) Expression of selected genes based on Reactome enrichment analysis (1I). RNA-sequencing shows that silencing of *Id1* leads to upregulation of genes associated with epidermal differentiation, whereas genes linked to adhesion and extracellular matrix are down regulated. Data are represented as mean ± SEM. ***p < 0.001 using multiple unpaired t-test. Scale bar 50 μm.

In addition to *Id1*, *Id2 and Id3* are also expressed in the developing epidermis. Expression analysis of *Id2* and *Id3* using E13 single-cell RNA-sequencing fails to reveal enrichment in committed progenitors (cluster 2) (Figure S1H and S1I). Analysis of *Id* coexpression shows that more than half (59%) of E13 epidermal progenitors express *Id1* but not *Id2* or *Id3*, whereas 36% of progenitor cells co-express all three *Id* members. Only a small percentage (5%) of progenitors at E13 are *Id* negative (Figure S1J). These data suggest that ID proteins likely have both redundant and non-redundant functions in epidermal development.

### ID1 is associated with epidermal lineage progression

To address the function of ID1 in epidermal progenitor cells, we transduced primary epidermal progenitor cells with either control scramble shRNA (*shScr*) or an shRNA targeting *Id1* (*shId1*) (Figure 1F-1G). In line with the distinct expression profiles of *Id1* compared to *Id2* and *Id3* (Figure S1H and S1I), silencing of *Id1* in cultured epidermal progenitors did not lead to a compensatory upregulation of *Id2* and *Id3* mRNA (Figure 1H). Transcriptional profiling revealed that differentially expressed genes (>2x fold change) were both up (76%) and down (24%) regulated upon *Id1* silencing (Figure S1K). Reactome pathway analysis including DEGs upregulated in *shId1* compared to *shScr* transduced primary epidermal progenitors revealed biological processes associated with formation of the cornified envelope and keratinization, indicative of epidermal differentiation. In contrast, down regulated DEGs were linked to extracellular matrix (ECM) modulation and laminin signaling (Figure 1I-K), suggesting that *Id1* influence gene programs altering the composition of the ECM and/or adhesion to the basement membrane. All in all, the *in vivo* ID1 expression pattern in combination with transcriptional profiling, associates ID1 with epidermal lineage progression.

### ID1 counteracts epidermal progenitor delamination

To assess the function of ID1 *in vivo*, we exploited the method of ultrasound guided *in utero* injections of high titer lentiviral particles (Beronja et al., 2010)(Figure 2A). *Id1* was either silenced in wild type mice using an shRNA targeting *Id1*, or depleted by injection of a CRE-expressing lentivirus (LV-CRE) into a conditional *Id1* transgenic (*Id1^fl/fl^* mouse line at E9.5 (Nam and Benezra, 2009). Delivery of *shId1* (as judged by H2BGFP reporter expression) successfully reduced *in vivo* ID1 protein expression at E14.5 when compared to embryos targeted with *shScr,* in line with *in vitro* knock down efficiency (Figure 2B and 2C). Similarly, LV-CRE injection resulted in efficient loss of ID1 immunoreactivity in *Id1*^fl/fl^ targeted epidermal progenitors at E14.5, whereas ID1 protein was sustained in *Id1*^+/fl^ targeted skin (Figure S2A), indicating that injection of either *shId1* or LV-CRE efficiently ablated epidermal ID1 expression *in vivo*.

**Figure 2.**
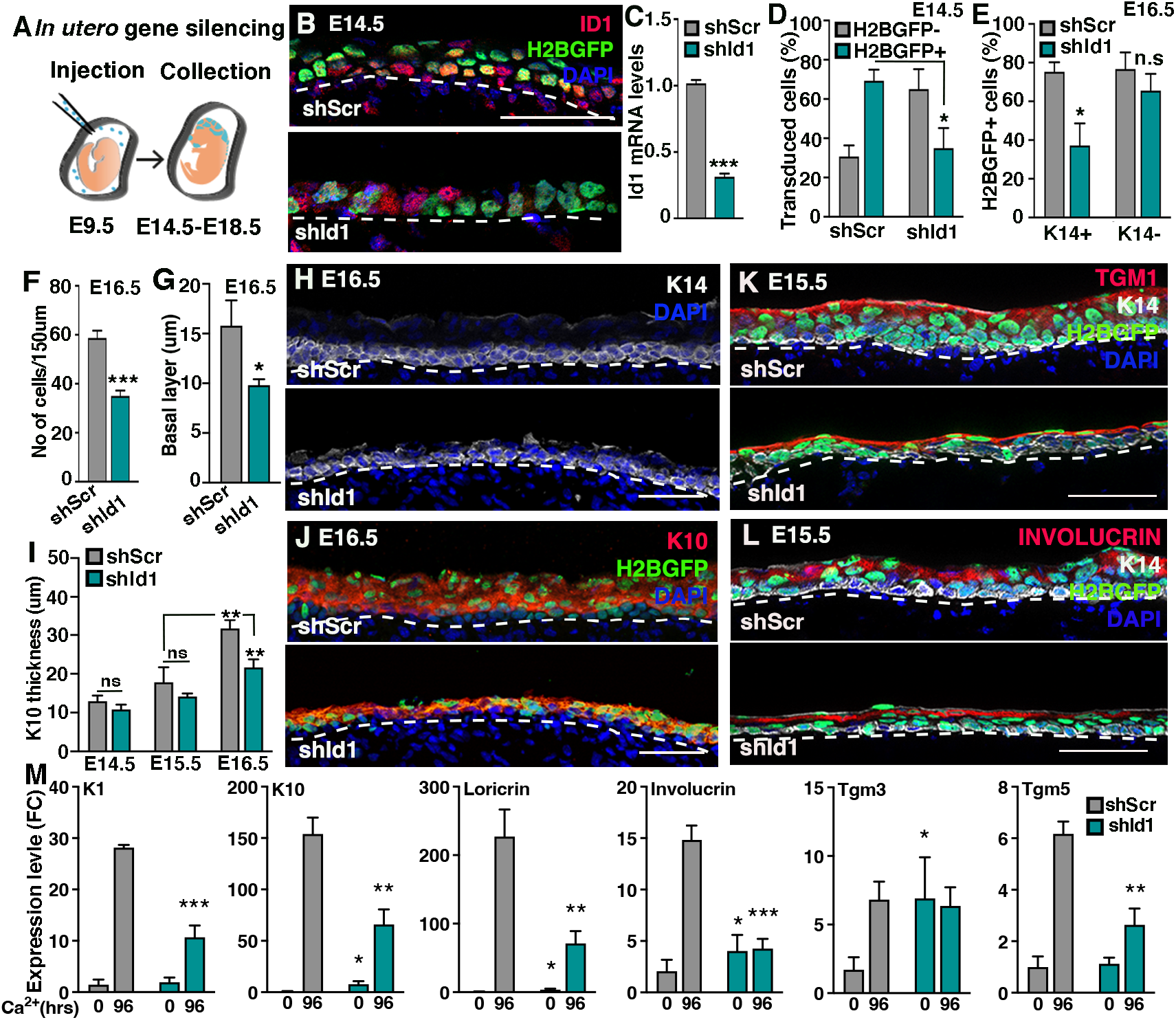
ID1 counteracts epidermal progenitor delamination. (A) Schematic representation of *in utero* lentiviral injections. Virus is injected into the amniotic cavity at E9.5 and embryos are collected between E14.5 and E18.5. (B) E14.5 ID1 immunoreactivity is reduced in H2BGFP reporter positive cells upon injection of lentiviral *shId1,* but not control *shScr.* (C) *Id1* mRNA expression is reduced in epidermal progenitors targeted *in vitro* with *shId1* compared to *shScr*. (D) The percentage of transduced H2BGFP-positive epidermal progenitors is reduced in *shId1* targeted epidermis compared to *shScr* at E14.5. (E) Quantification of percentage of H2BGFP positive targeted K14 expressing basal progenitors and H2BGFP positive, K14 negative suprabasal keratinocytes indicate selective loss of *shId1* targeted progenitor cells. (F) Total number of epidermal nuclei per 150um is significantly reduced in *shId1* compared to *shScr* targeted epidermis at E16.5. (G) Quantification of K14-positive basal progenitor layer thickness in *shId1* and *shScr* epidermis at E16.5. (H) Representative images of K14 thickness at E16.5 in *shId1* and *shScr* epidermis. (I) K10-positive spinous layer thickens with epidermal development. Silencing of *Id1* impairs spinous layer development. (J) Representative images of K10 distribution at E16.5 in *shId1* and *shScr* epidermis. (K and L) Terminal differentiation is temporally and spatially normal in *shId1* compared to *shScr* as judged by expression of differentiation marker TGM1 and INVOLUCRIN at E15.5. (M) Differentiation markers expression in *shScr* and *shId1* cells. Statistical analysis is performed between *shScr* and *shId1* within 0 or 96 hrs, respectively. Data are represented as mean ± SD. *p < 0.05, **p < 0.01, ***p < 0.001 using multiple unpaired t test. Scale bars 50 μm.

Even though the transduction efficiency of *shId1* and *shScr* were comparable when cultured epidermal progenitors were infected (Figure S2B), the percentage of H2BGFP reporter-expressing progenitors was reduced in E14.5 *shId1* epidermis compared to *shScr* (Figure 2D and S2C). At E16.5, *shId1* embryos displayed significantly fewer H2BGFP/K14 positive progenitor cells compared to embryos targeted with *shScr*, whereas the number of H2BGFP positive, K14 negative suprabasal cells remained high (Figure 2E and S2D). At E18.5, *shId1* targeted basal progenitor cells were scarce and remaining epidermal H2BGFP expression was localized to K14 negative, suprabasal keratinocytes or developing hair follicles (Figure S2E). Analogously, we found a reduction of the number of *Id1* silenced progenitor cells at E16.5 in *Id1^fl/fl^* embryos when compared to E14.5 (Figure S2F-S2I), indicating that progenitors lacking ID1 are selectively lost during epidermal development.

To assess if loss of *Id1* targeted progenitors is compensated by untargeted progenitors, we quantified the total number of cells in *shId1* epidermis at E14.5 and E16.5. We found a reduction in the epidermal cell number at E16.5, but not E14.5, in *shId1* epidermis compared to *shScr* targeted embryos (Figure 2F and S2J). We then analyzed apoptosis to see if cell death could explain the selective loss of *Id1* depleted progenitors but did not detect increased CC3-immunoreactivity in *shId1* compared to *shScr* targeted E14.5 epidermis (Figure S2K and S2L). Collectively, these results suggest that ID1 counteracts precocious epidermal progenitor cell delamination, independent of apoptosis.

### Loss of ID1 results in epidermal thinning without affecting terminal differentiation

To assess if the reduction of epidermal cell number at E16.5 corresponded to loss of progenitor or differentiated cell layers, we first quantified the thickness of the entire epidermis as well as the basal K14-positive progenitor layer and found both to be diminished when comparing *shId1* to *shScr* embryos (Figure 2G-H and S2M). Since our *in vitro* transcriptional profiling implicated ID1 in epidermal differentiation (Figure 11 and 1J), we characterized the dynamic formation of the K10 expressing spinous layer between E14.5 to E16.5. Whereas the thickness of the spinous layer increased in *shScr* targeted epidermis during development, it failed to do so efficiently in *shId1* infected epidermis (Figure 2I), and at E16.5 there was a significant reduction of the thickness of the spinous layer in *shId1* epidermis when compared to *shScr* targeted epidermis (Figure 2I and 2J). Due to the rapid loss and few remaining CRE targeted cells in *Id1^fl/fl^* epidermis (Figure S2H), we failed to detect any reduction in either the E16.5 spinous layer, or the overall epidermal thickness (Figure S2N-P) suggesting that the expanding epidermis is able to compensate for the mosaic genetic depletion of ID1.

*In vivo* ID1 expression pattern is consistent with a role for ID1 in commitment to differentiation rather than terminal differentiation. To assess if the absence of ID1 induced precocious terminal differentiation *in vivo,* we analyzed E15.5 epidermis, the earliest time point where we could confidently detect markers of terminal differentiation. Although we found a general thinning of the spinous and granular differentiated cell layers marked by TGM1 and INVOLUCRIN in *shId1* epidermis compared to *shScr* transduced skin (Figure 2K and 2L), TGM1 and INVOLUCRIN were correctly expressed, temporally as well as spatially. Cultured *shId1* and *shScr* progenitor cells both upregulated spinous and granular differentiation markers when asked to terminally differentiate for 96 hours *in vitro* (Figure 2M), although the upregulation was less prominent when *Id1* was silenced. Taken together, these data suggest that progenitor cells lacking ID1 do not initiate terminal differentiation precociously.

### Progenitor cells devoid of ID1 co-express basal and differentiation markers

During epidermal development, basal progenitor cell crowding induces delamination and subsequent differentiation (Miroshnikova et al., 2018). Delaminating epidermal progenitors couple induction of spinous fate markers with suppression of basal gene programs, and failure to do so leads to aberrant stratification (Blanpain et al., 2006). Revisiting our transcriptomics analysis (Figure 1I), we identified upregulation of spinous markers in *shId1* targeted epidermal progenitors *in vitro* (Figure 3A), suggesting that the basal-to-spinous transition is affected in the absence of ID1. Analysis of basal (K5) and spinous (K10) marker co-expression in E16.5 epidermis revealed enrichment of K5/K10 double-positive epidermal progenitors in *shId1* compared to *shScr* (Figure 3B and 3C). Analogously, the percentage of K5/K10 double-positive progenitors was significantly increased in LV-CRE targeted *Id1^fl/fl^* compared to *Id1^+/fl^* epidermis (Figure 3D and S3A), indicating that loss of ID1 leads to an accumulation of cells co-expressing markers normally representative of distinct cell states.

**Figure 3.**
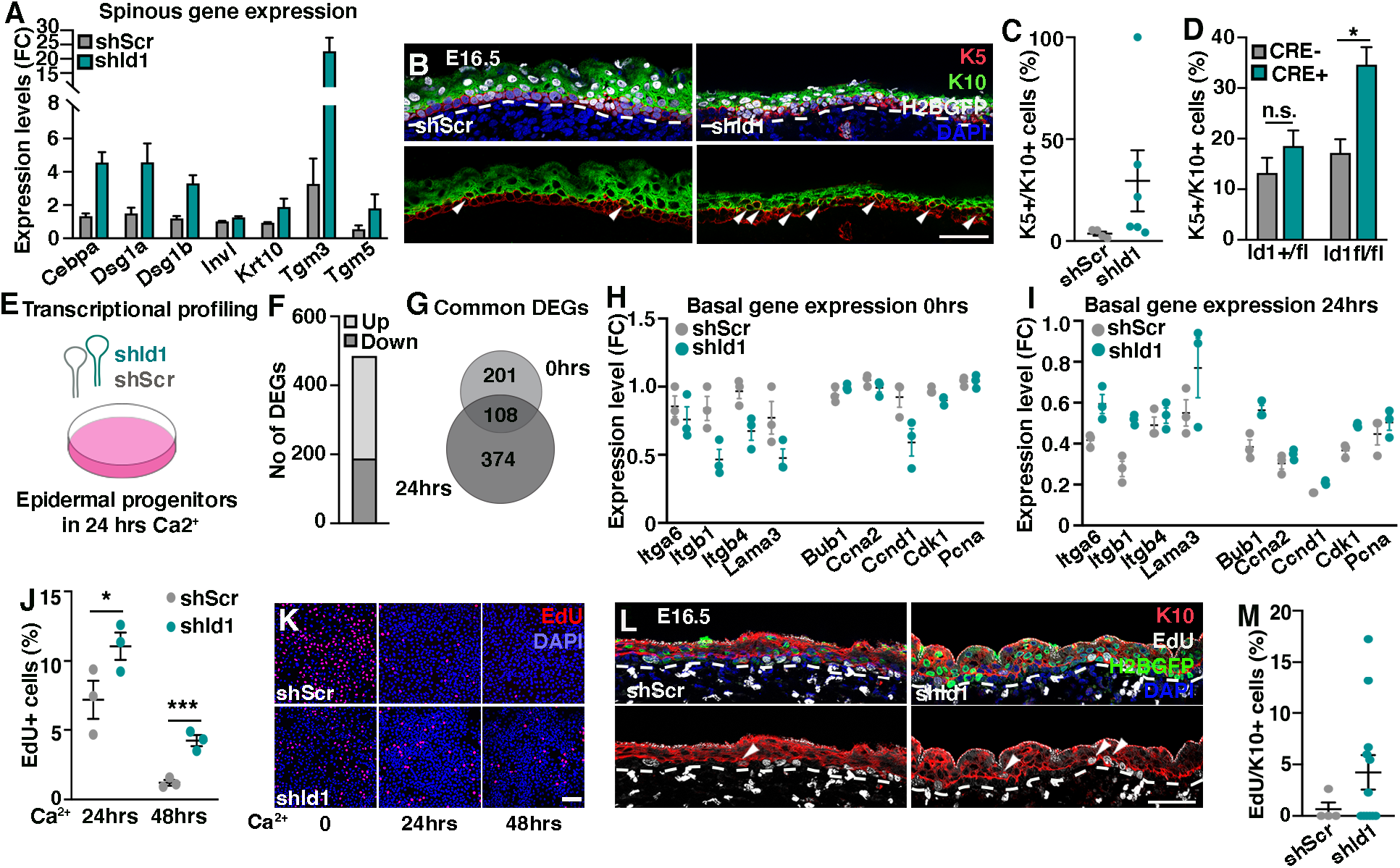
Progenitor cells devoid of ID1 co-express basal and differentiation markers. (A) Spinous marker expression after silencing of *Id1* in cultured epidermal progenitor cells. (B and C) *In vivo* silencing of *Id1* increases the incidence of K5/K10 double-positive cells at E16.5 compared to *shScr* targeted epidermis. (D) The percentage of K5/K10 double-positive cells is significantly increased in *Id1^fl/fl^* compared to *Id1^+/fl^* epidermis when ID1 is ablated using a CRE lentivirus. (E) Cultured epidermal progenitors were targeted with *shId1* or *shScr* and asked to differentiate for 24 hours before collection and transcriptional profiling. (F) Number of differentially expressed genes in *shId1* compared to *shScr* targeted cells at 24 hours of differentiation. (G) Overlap between differentially expressed genes at 0 and 24 hours of differentiation suggest that ID1 has overlapping and distinct functions in epidermal lineage progression. (H) Expression of genes associated with a basal progenitor cell state are reduced in *shId1* compared to *shScr* targeted cultured epidermal progenitor cells. (I) Basal genes are sustained in *shId1* targeted progenitors upon differentiation compared to control *shScr* targeted cells. (J and K) EdU incorporation upon differentiation of *shId1* and *shScr* infected progenitors. Scale bar 100 μm. (L and M) *In vivo* silencing of *Id1* leads to an increased number of suprabasal cells coexpressing EdU and K10 when compared to control epidermis. Data are represented as mean ± SEM. *p < 0.05, ***p < 0.001 using multiple unpaired t-test. Scale bars 50 μm if not specified.

### ID1 co-ordinate the basal to suprabasal progenitor state transition

Interestingly, most cells with double progenitor/differentiation marker expression were not in contact with the basement membrane (Figure 3B), suggesting that ID1 silenced progenitors failed to downregulate progenitor markers during delamination. To this aim, we sequenced *shId1* and *shScr* targeted epidermal progenitors after 24 hours of *in vitro* differentiation (Figure 3E). In line with transcriptional alterations found in *shId1* targeted epidermal progenitors, differentially expressed genes (>2x fold change) were both up (61%) and down (39%) regulated. Furthermore, Reactome analysis confirmed the involvement of ID1 in coupling inhibition of differentiation to modulation of extracellular matrix (Figure 3F and S3B). Interestingly, 35% of the differentially expressed genes found in *shId1* targeted epidermal progenitors were significantly altered upon differentiation of *shId1* progenitors (Figure 3G) suggesting that aspects of ID1 function are conserved in basal progenitors and differentiating keratinocytes.

Focusing on markers associated with the basal state revealed that silencing of *Id1* in progenitor cells resulted in reduced expression of genes linked to basal membrane anchoring and cell cycle progression (Figure 3H), potentially coupling loss of ID1 to progenitor cell delamination and cell cycle exit. Paradoxically however, comparing transcriptional profiles of *shId1* and *shScr* progenitors after 24 hours of differentiation revealed that *shId1* targeted progenitors sustained expression of basal markers normally lost during differentiation (Figure 3I). These data suggest that although epidermal progenitors down regulate basal markers upon ID1 silencing, their response to *in vitro* differentiation cues is impaired.

To address if silencing of *Id1* not only affects expression of a cohort of basal genes, but also affects basal cell characteristics during lineage progression, we pulsed *in vitro* differentiated epidermal progenitors with EdU. Whereas as expected, control *shScr* progenitors rapidly exited the cell cycle and were largely EdU negative already at 24 hours of differentiation, a significant fraction of differentiated progenitors targeted with *shId1* still incorporated EdU after 48 hours of differentiation (Figure 3J and 3K). In addition, we observed an enrichment of suprabasal EdU/K10 double-positive progenitors in *shId1* silenced epidermis normally absent in *shScr* targeted epidermis at E16.5 (Figure 3L and 3M). Our work suggests that ID1 impinges on the basal to suprabasal transition, acting to co-ordinate and fine-tune progenitor as well as differentiation gene programs throughout epidermal lineage transition.

### Epidermal progenitor proliferation is positively regulated by ID1

Tissue development requires regulation of mechanisms balancing progenitor cell proliferation and differentiation. Since down regulation of ID1 leads to diminished expression of basal progenitor cell markers (Figure 3H), including the cell cycle regulator *Cnnd1*, we analyzed epidermal progenitor proliferation after manipulation of *Id1. In vivo* targeting of *Id1* reduced progenitor cell proliferation using either an shRNA or LV-CRE strategy (Figure 4A-B and S4A), and cultured primary *shId1* epidermal progenitor cells incorporated significantly less EdU than *shScr* progenitors (Figure 4C and 4D). In contrast, overexpression of ID1 in primary epidermal progenitor cells increased EdU incorporation *in vitro* (Figure 4E-F and S4B-S4C), suggesting that ID1 positively regulate progenitor renewal during epidermal development.

**Figure 4.**
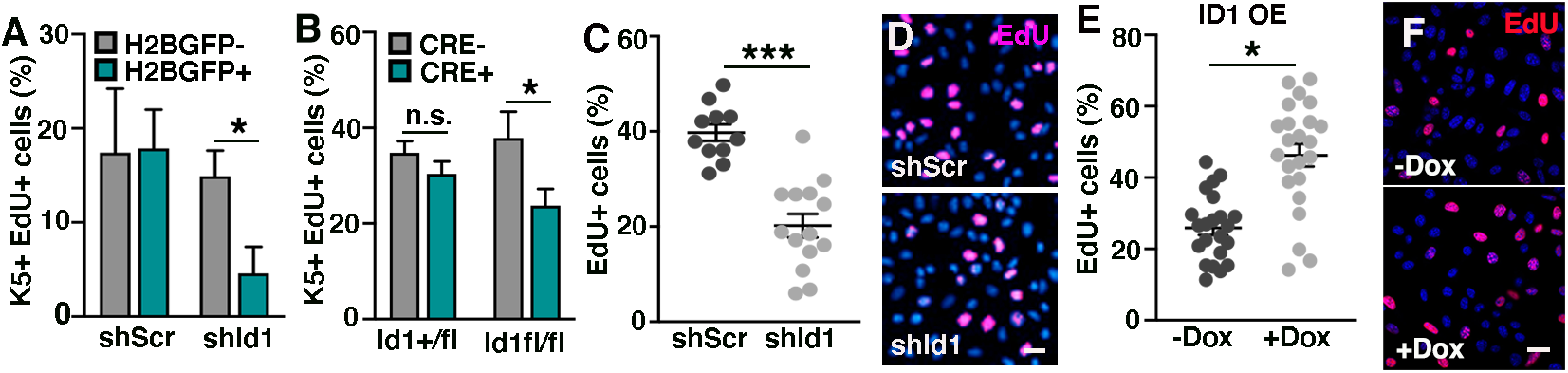
Epidermal progenitor proliferation is positively regulated by ID1. (A) Percentage of *in vivo* K5/EdU-positive proliferating progenitors are reduced upon *shId1* targeting compared to *shScr* targeted epidermis at E16.5. (B) LV-CRE mediated *Id1* ablation reduces proliferation in K5 positive epidermal progenitors in *Id1^fl/fl^* compared to *Id1^+/fl^* epidermis. (C and D) Silencing of *Id1* in cultured epidermal progenitor cells leads to reduced EdU incorporation. (E and F) Overexpression of ID1 in cultured epidermal progenitors leads to increased EdU incorporation. Data are represented as mean ± SEM. *p < 0.05, ***p < 0.001 using multiple unpaired t-test. Scale bars 25 μm.

### Identification of ID1 gene signatures

Our data positions ID1 at the intersection of proliferation and differentiation in epidermal progenitor cells. Reasoning that silencing of *Id1* negatively affects proliferation and leads to an induction of differentiation markers, forced ID1 expression would, in addition to driving cell cycle progression, shift gene expression profiles towards a basal state. To this end, we profiled epidermal progenitor cells overexpressing ID1, as well as ID1 overexpressing progenitors asked to differentiate for 24 hours (Figure 5A and S5A). Focusing on down regulated DEGs (>2x fold change) at 24 hours of differentiation, GO analysis revealed significantly enriched biological processes such as epidermal progenitor cell differentiation and regulation of signaling (Figure 5B), suggesting that differentiation is retarded when ID1 is overexpressed. Expression levels of spinous markers elevated in *shId1* progenitors, were in contrast found to be reduced in ID1 overexpressing progenitors when compared to uninduced progenitors, albeit from a low starting level (Figure 5C). After 24 hours of differentiation, induction of spinous gene expression in ID1 overexpressing cells was prominent, although hampered, when compared to uninduced differentiated cell (Figure S5B). In contrast to cells silenced for ID1, we did not find alterations in basal gene expression (Figure S5C and S5D). These data suggest that sustained ID1 expression during differentiation acts to slow down, rather than inhibit epidermal differentiation.

**Figure 5.**
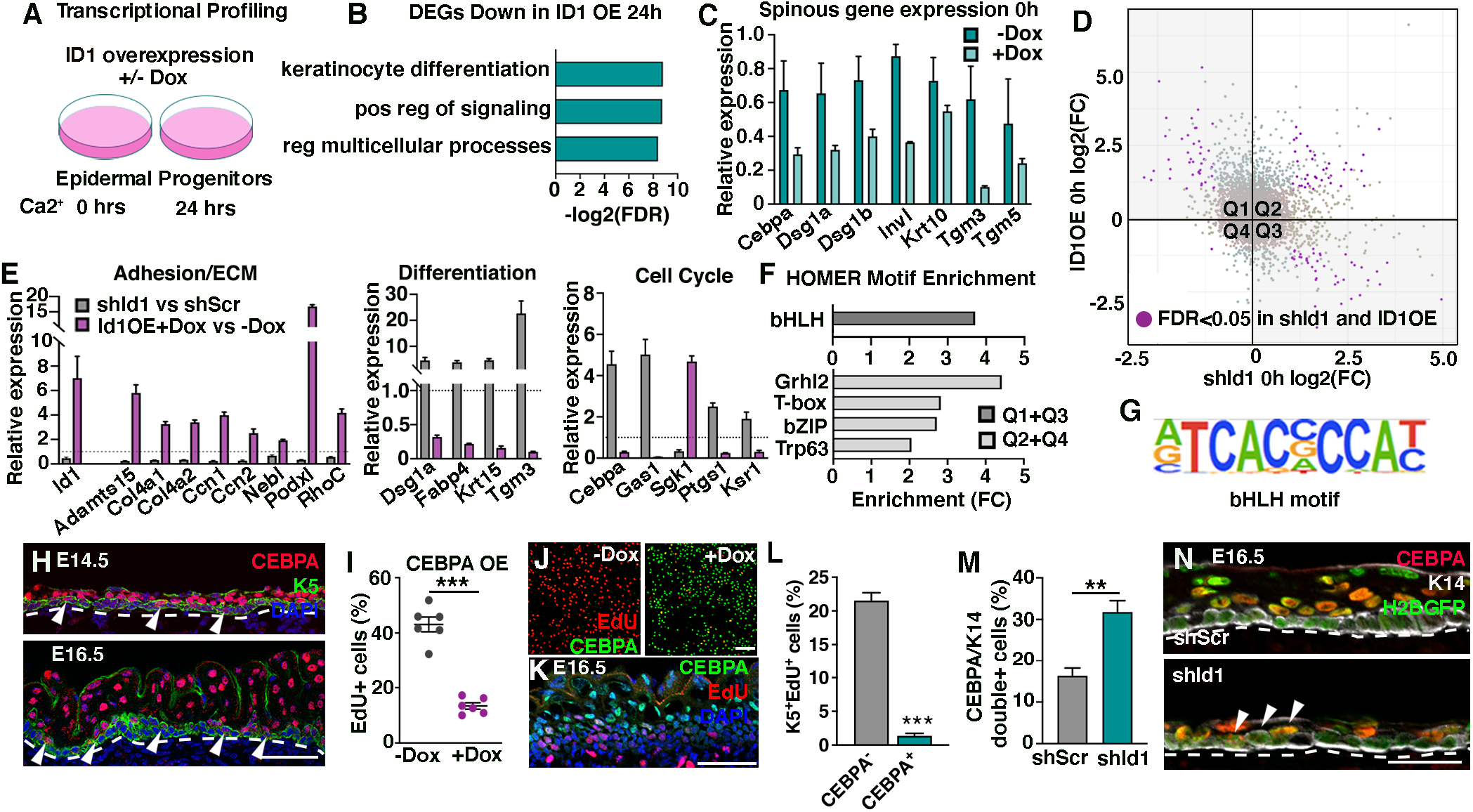
Identification of ID1 gene signatures. (A) ID1 is induced in a doxycycline dependent manner and progenitor cells are differentiated and transcriptionally profiled. (B) Downregulated genes after ID1 overexpression and 24 hours of differentiation are linked to keratinocyte differentiation and regulation of signaling. (C) Spinous gene expression is reduced in epidermal progenitors upon ID1 overexpression. (D) Identification of ID1 gene signatures by merging expression data from *shId1* and ID1 overexpression profiling of epidermal progenitors reveal a cohort of statistically significant genes (marked in purple) (LogFC >1 and FDR >0.05). Q1 and Q3 represent genes whose expression correlates to modulation of ID1. Q2 and Q4 represent genes that are only induced (Q2) or repressed (Q4) when ID1 levels are altered. (E) ID1 signature genes are functionally linked to adhesion and extracellular matrix modulation, differentiation or cell cycle regulation. (F) HOMER transcription factor binding motif analysis show distinct enrichment binding motifs in promoters (+400 bp) of genes found in Q1+Q3 compared to promoters in the expressed transcriptome. (G) bHLH motif enriched in Q1+Q3 gene promoters. (H) Immunoreactivity of CEBPA and K5 in E14.5 and E16.5 localizes CEBPA to suprabasal keratinocytes and scattered basal K5-positive progenitors. Arrows indicate CEBPA-positive basal cells. (I and J) Overexpression of CEBPA significantly reduces EdU incorporation *in vitro*. Quantification done after 3 days of doxycycline treatment. Scale bar 100 μm. (K and L) Proliferating K5-positive progenitors are largely CEBPA negative. Data are presented as mean ± SD. (M and N) K14/CEBPA double-positive progenitors are significantly enriched in *shId1* targeted epidermis when compared to *shScr*. Arrows highlight K14/CEBPA positive cells. Data are represented as mean ± SEM. **p < 0.01 ***p < 0.001 using multiple unpaired t-test. Scale bars 50 μm except for J (100 μm).

Combining transcriptional profiling results from both *shId1* and ID1 overexpression in epidermal progenitor cells allowed us to define a set of 83 candidate *Id1* target genes whose expression changes correlated significantly (FDR>0.05) with alteration of *Id1* transcripts levels (Figure 5D, Q1 and Q3, Table S1). Within this relatively small group of candidates, we identified cohorts of genes representing aspects of the *in vivo* phenotypes described upon *Id1* silencing (Figure 5E), correlating alterations in *Id1* associated gene signatures to ECM modulation, differentiation and cell cycle regulation respectively.

### ID1 interacts directly with bHLH transcription factors

Lacking a DNA binding domain, ID1 is unable to affect transcription through direct chromatin interaction at target genes (Massari and Murre, 2000). To mechanistically begin to understand how ID1 affects epidermal progenitor cells, we aimed to identify epidermal ID1 binding partners. ID1, as well as family members ID2 and ID3, were overexpressed in cultured epidermal progenitor cells, after which ID-binding protein complexes were isolated and analyzed using mass spectrometry (Figure S5E and S5F). We identified three known class I bHLH transcription factors, TCF3 (E2A), TCF4 (ITF2) and TCF12 (HEB) to be bound to ID1, whereas ID2 and ID3 associated exclusively with TCF12 (Table S2). Other published ID1-interacting transcription factors (Roberts et al., 2001; Yates et al., 1999) were not identified, suggesting that ID1 predominantly affects gene expression in the epidermis through binding to the TCF bHLHs. Generally expressed class I bHLH transcription factors, such as TCF3/4/12, often heterodimerize with cell type specific class II bHLH factors (Murre et al., 1989a; Murre et al., 1989b), thereby acquiring cell state and context specific target gene profiles. Considering that TCF3 and TCF4 exclusively interacted with ID1 and no other ID family member, we focused on identifying TCF3/4 heterodimerizing partners in addition to ID1. Expression and purification of TCF3 and TCF4 in primary epidermal progenitor cells however failed to identify additional bHLH interactors but could independently confirm the binding of TCF3 and TCF4 to ID1 respectively (Table S2). Collectively, these data suggest that ID1 binding to TCF3, TCF4 and TCF12 acts to modulate bHLH transcriptional programs during epidermal development.

Transcription factor binding motif analysis on promoter sequences in genes differentially expressed in *shId1* and ID1 OE (Figure 5D, Q1 and Q3) using HOMER motif discovery revealed enrichment of bHLH motifs in potential ID1 targets when compared to all genes expressed epidermal progenitors (Figure 5F and 5G). Interestingly, motif discovery in genes significantly altered, but uncorrelated to ID1 levels (Figure 5D, Q2 and Q4), revealed distinct motif profiles highlighting known regulators of epidermal differentiation such as GRHL, bZIP (basic leucine zipper domain), T-box and Trp63 transcription factors (Figure 5F and S5G). Taken together, these data suggest that bHLH transcriptional programs impact epidermal progenitor states through modulation of specific target genes.

### CEBPA is ectopically expressed in epidermal progenitor cells in the absence of ID1

Having identified an *in vitro* ID1 gene signature harboring enrichment of bHLH binding motifs, we singled out CEBPA (CCAAT Enhancer Binding Protein Alpha) for further in-depth characterization. CEBPA is an interesting ID1 signature gene - the combined loss of CEBPA and CEBPB in the developing epidermis results in hyperproliferation and epidermal barrier defects (Lopez et al., 2009), largely resembling silencing of *Id1.* We found prominent CEBPA expression in suprabasal layers at E14.5 and E16.5. In addition, a significant subset of K5-positive basal progenitors expressed lower levels of CEBPA (Figure 5H). *In vitro* differentiation of epidermal progenitor cells mimicked the *in vivo* expression pattern with significant upregulation of *Cebpa* mRNA and protein concomitant with differentiation (Figure S5H and S5I). In line with previous reports (Lopez et al., 2009), we found that doxycycline-mediated overexpression of CEBPA in cultured epidermal progenitors significantly reduced the number of EdU^+^ cycling progenitor cells (Figure 5I-J and S5J). Quantification of EdU-incorporation in the K5-positive basal layer at E16.5 reveals that most proliferating cells are CEBPA negative (Figure 5K-L), confirming a role for CEBPA in repressing epidermal progenitor cells proliferation. Supporting our *in vitro* sequencing data, shRNA silencing of *Id1* resulted in an increase in the number of K14/CEBPA double-positive progenitor cells compared to *shScr* at E16.5 (Figure 5M-N) indicating that CEBPA expression is repressed in the presence of ID1 *in vivo*. To mechanistically explore if *Cebpa* is transcriptionally regulated through an ID1-TCF axis, we cloned a 2kb fragment of the *Cebpa* promoter as well as an upstream *Cebpa* enhancer (Cooper et al., 2015) containing reported functional bHLH binding E-box motifs (Cooper et al., 2015; Hu et al., 2016; Pfurr et al., 2017; Soleimani et al., 2012; Yoon et al., 2015). We found an increase in promoter and enhancer luciferase reporter activity in *shId1* epidermal progenitor cells when compared to *shScr* (Figure S5K-L) correlating to the observed increase in mRNA and *in vivo* protein expression, suggesting that *Cebpa* regulatory elements are differentially engaged in the presence or absence of ID1.

### TCF3/4/12 localize to the developing epidermis and regulate progenitor cell proliferation

TCF3, TCF4 and TCF12 are established class I bHLH ID interactors (Lasorella et al., 2001), but the expression dynamics and function of TCFs in the developing epidermis has not been elucidated. Returning to the E13 epidermal single-cell profiling (Fan et al., 2018), we found all three *Tcfs* to be ubiquitously expressed in a majority of epidermal progenitor cells in both cluster 1 and 2, contrasting the cluster 1 enrichment of *Id1* (Figure S6A-B). *In situ* hybridization confirmed the expression of *Tcf3, Tcf4* and *Tcf12* mRNA in epidermal progenitors at E14.5 (Figure 6A). Later in epidermal development (E18.5), *Tcf3, Tcf4 and Tcf12* mRNA reactivity became enriched in basal progenitors compared to suprabasal differentiated keratinocytes (Figure 6A). We also confirmed high expression of all three *Tcfs* in the dermis at the timepoints analyzed (Rezza et al., 2016; Sennett et al., 2015). In line with the E13 single-cell analysis, *in vitro* differentiation of cultured epidermal progenitors did not distinctively alter the mRNA expression of either *Tcf3, Tcf4 or Tcf12* (Figure S6C), supporting a relatively broad expression of bHLH TCFs in the developing epidermis. Our data support a model where the bHLH transcriptional output is controlled by spatially restricting ID protein expression.

**Figure 6.**
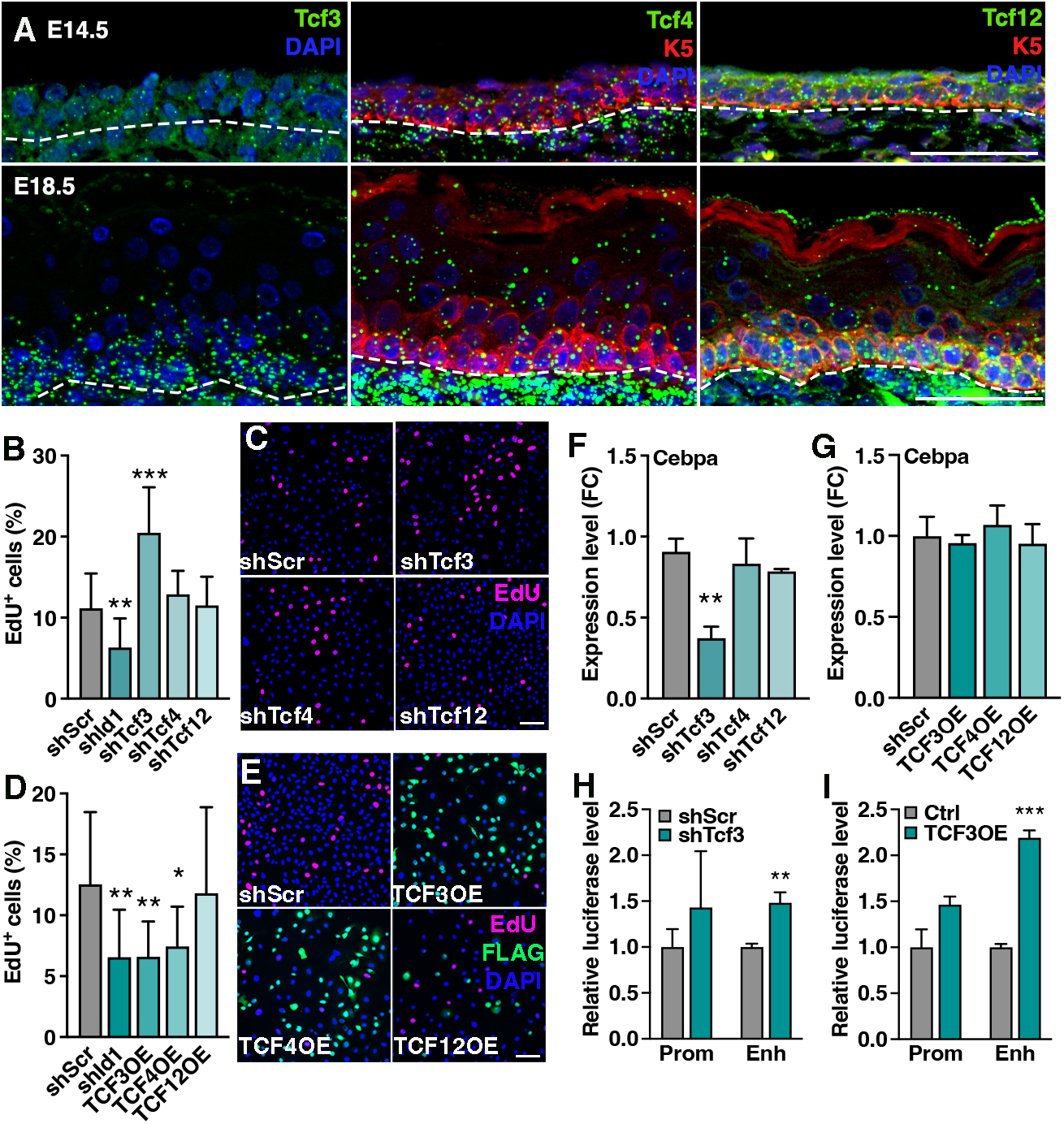
TCF3/4/12 localize to the developing epidermis and regulate progenitor cell proliferation. (A) Visualization of *Tcf3, Tcf4* and *Tcf12* mRNA in skin reveal expression in epidermal progenitors at E14.5, with subsequent enrichment in basal layer upon induction of differentiation (E18.5). Scale bar 50 μm. (B) Silencing of *Tcf3*, but not *Tcf4* or *Tcf12* promotes progenitor cell proliferation. (C) Representative images showing EdU incorporation in keratinocytes targeted with *shScr, shTcf3, shTcf4,* and *shTcf12.* Scale bar 100 μm. (D) Overexpression of TCF3 and TCF4 reduces proliferation *in vitro*. (E) Representative images showing EdU incorporation in keratinocytes overexpressing TCF3, TCF4, and TCF12 (transient overexpression in *shScr* keratinocytes). Scale bar 100 μm. (F) Silencing of *Tcf*3 reduces *Cebpa* mRNA levels. (G) *Cebpa* mRNA is not altered upon forced TCF expression. (H-I) Cebpa promoter and enhancer luciferase activity upon Tcf3 silencing (H) and overexpression (I). Data are represented as mean ± SD. *p < 0.05 **p < 0.01 ***p < 0.001 using multiple unpaired t-test.

To address the function of bHLH TCF transcription effectors in epidermal progenitor cells, we silenced *Tcf3, Tcf4* and *Tcf12* and assessed proliferation. Knock down of *Tcf3,* but not *Tcf4* or *Tcf12*, resulted in increased EdU incorporation *in vitro* (Figure 6B-C and S6D). In contrast, overexpression of TCF3 and TCF4, but not TCF12 reduced proliferation in cultured epidermal progenitor cells (Figure 6D-E), suggesting that ID1 regulates epidermal progenitor cell proliferation through TCF3-, and possibly TCF4-, dependent transcriptional programs.

### *Cebpa* expression is bHLH-independent in epidermal progenitors

Our data suggest that activation of CEBPA is linked to silencing of *Id1*, *in vitro* and *in vivo*. To address if ID1 acts to suppress *Cebpa* gene expression through sequestering of bHLH TCF transcriptional effectors, we assessed *Cebpa* mRNA expression after silencing or overexpression of TCF3, TCF4 and TCF12 (Figure 6F-G). We failed to see upregulation of *Cebpa* mRNA when TCFs were overexpressed, however silencing of *Tcf3* reduced *Cebpa* levels. Focusing on TCF3, we found that the *Cebpa* promoter and enhancer activity did not correlate to TCF3 levels (Figure 6H-I), arguing that bHLH TCF factors are not required for *Cebpa* transcription. Since our data suggest that upregulation of *Cebpa,* a known marker of epidermal differentiation, is independent of TCFs, we asked if silencing of *Tcf3* affects other genes associated with differentiation. Interestingly, epidermal progenitor cells targeted with *shTcf3,* but not consistently *shTcf4* or *shTcf12,* downregulated markers associated with differentiation which were found upregulated in *shId1* targeted progenitors (Figure S6E). Taken together, these data suggest that whilst CEBPA expression negatively correlates with ID1, *Cebpa* is not a direct bHLH target gene in epidermal progenitor cells.

### pSMAD1/5 activation of the *Id1* promoter is CEBPA dependent

We noticed that modulating CEBPA expression affects *Id1* mRNA and protein (Figure 7A-B and S7A), where high CEBPA expression acts to suppress ID1. *Id1* is a bona fide BMP target in the hair follicle stem cell lineage (Genander et al., 2014), and BMP signaling, as judged by phosphorylation of downstream effectors SMAD1/5, is active in epidermal progenitors and during *in vitro* differentiation (Figure 7C). Treatment of epidermal progenitor cells with BMP4 for 3 hours in culture induced *Id1* mRNA in both proliferating progenitors and keratinocytes differentiated for 24 hours (Figure 7D). pSMAD1/5 binding to the *Id1* promoter is well characterized (Genander et al., 2014; Korchynskyi and ten Dijke, 2002), and we assessed BMP sensitivity in epidermal progenitors and differentiated keratinocytes using a BMP-responsive pSMAD1/5 binding region located distally in the *Id1* promoter (Genander et al., 2014; Korchynskyi and ten Dijke, 2002). Whereas the pSMAD1/5 binding region in the *Id1* promoter responded to BMP in epidermal progenitor cells, keratinocytes differentiated for 24 hours failed to activate *Id1* luciferase reporter activity in the presence of BMP (Figure 7E), suggesting that pSMAD1/5 engagement with the *Id1* promoter is dynamically regulated during epidermal differentiation.

**Figure 7.**
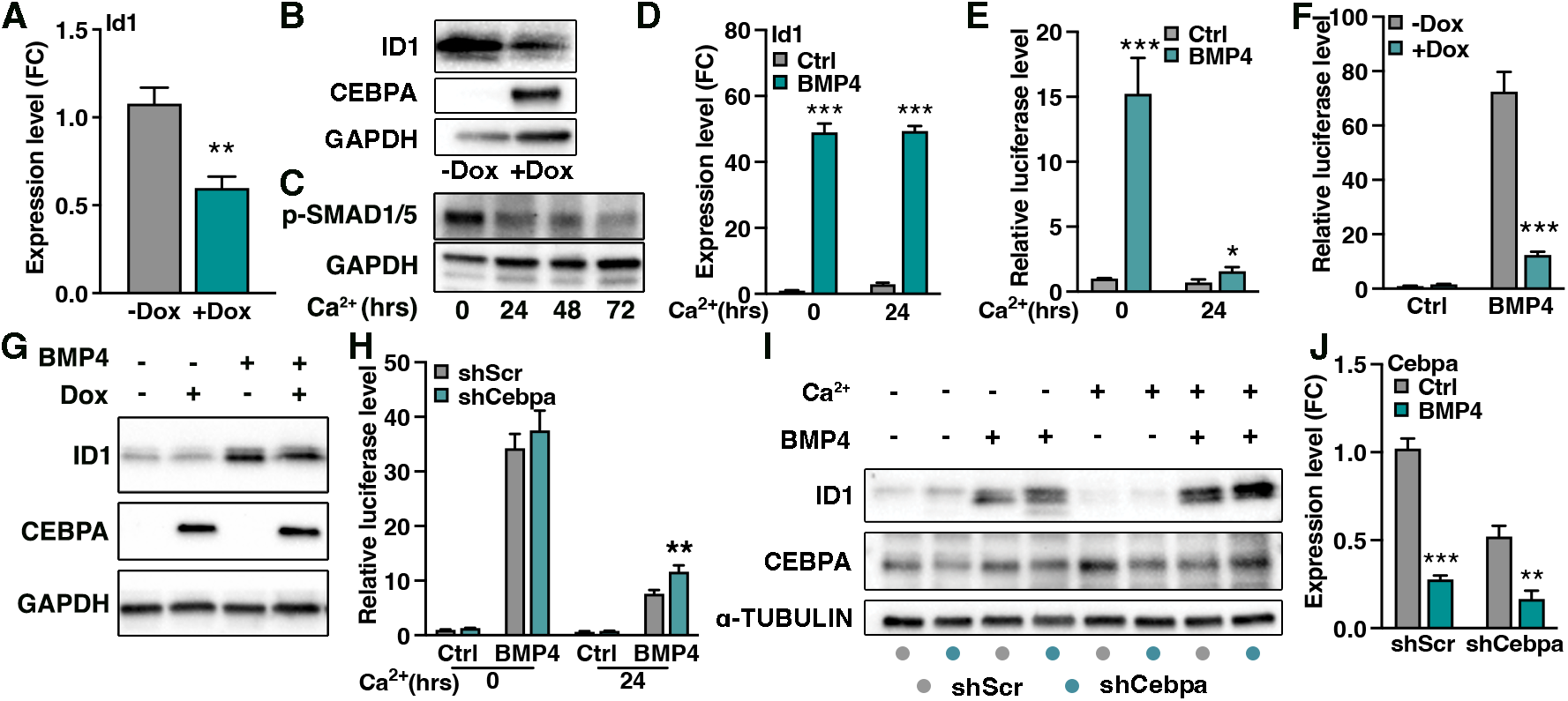
pSMAD1/5 activation of the *Id1* promoter is CEBPA dependent. (A) Relative *Id1* mRNA level upon 3 days of CEBPA overexpression. (B) Protein level of ID1 is reduced upon forced CEBPA expression. (C) pSMAD1/5 activity is pronounced in cultured epidermal progenitors and diminished upon differentiation. (D) Cultured epidermal progenitors (0 hrs Ca^2+^) and differentiated keratinocytes (24 hrs Ca^2+^) both respond to 3 hours BMP4 treatment by upregulation of *Id1* mRNA. (E) Luciferase reporter activity show pronounced *Id1* promoter activity in response to BMP in progenitors when compared to differentiated keratinocytes. (F) Forced CEBPA expression in epidermal progenitor cells inhibits the BMP-mediated activity of the *Id1* promoter. (G) Induction of ID1 is impaired after BMP4 treatment in the presence of CEBPA. (H) Silencing of *Cebpa* enhances *Id1* promoter activity in differentiated keratinocytes and leads to upregulation of CEBPA (I). (J) BMP4 treatment for 3 hours repress *Cebpa* expression. Data are represented as mean ± SD. *p < 0.05 **p < 0.01 ***p < 0.001 using multiple unpaired t-test.

CEBPA can redirect chromatin binding of SMADs during lineage commitment (Trompouki et al., 2011). To this end, we overexpressed CEBPA in epidermal progenitor cells and found a dramatic reduction in BMP-induced response of the *Id1* promoter, which correlated with reduced induction of ID1 protein levels (Figure 7F-G). To ask if CEBPA normally acts to repress pSMAD1/5-dependent *Id1* expression, we silenced *Cebpa* in differentiated keratinocytes, a cell state where CEBPA levels are endogenously high (Figure 5). We found that down regulation of CEBPA potentiated BMP-mediated pSMAD1/5 activation of the *Id1* promoter in differentiated keratinocytes, albeit at a low level (Figure 7H-I and S7B-D). Interestingly, whereas BMP treatment induces *Id1* mRNA (Figure 7D), we found a BMP-mediated repression of *Cebpa* (Figure 7J). Collectively these results suggest that while BMP can act upstream of both *Id1* and *Cebpa* effectively balancing their expression, CEBPA is able to desensitize the *Id1* promoter elements to inductive BMP-mediated pSMAD1/5 transcription during epidermal commitment to differentiation.

## Discussion

The transcriptional networks underlying progenitor differentiation in developing tissues are beginning to clear as more regulatory circuits are being identified. Here we characterize transcriptional regulators in the embryonic skin to identify a new role for ID1 in epidermal development and place ID1 in a context of previously known epidermal effectors. We exploit developmental transitions forming the stratified epidermis and identify transcriptional crosstalk impacting progenitor cell states during epidermal development. Using published transcriptomics describing E13 epidermis (Fan et al., 2018), we identify *Id1* to be associated with progenitor cells committing to differentiation concurrent with stratification. *Id* genes are previously implicated in human skin differentiation (Lopez-Pajares et al., 2015; Rotzer et al., 2006) and ID proteins are deregulated in human skin disease with impaired differentiation (Bjorntorp et al., 2003). Using *in utero* gene manipulation (Beronja et al., 2010), which in contrast to transgenic recombination strategies (Andl et al., 2004; Vasioukhin et al., 1999), allow for targeting of the uncommitted single-layered epidermal progenitor population, we observe that ID1 promotes proliferative self-renewal (Figure 4) and restricts commitment to differentiation during epidermal development *in vivo* (Figure 3).

We find that progenitor cells devoid of ID1 are lost upon stratification (Figure 2). Previous reports identified proliferation of basal progenitor cells as an inducer of epidermal stratification during development (Miroshnikova et al., 2018) where progenitor division result in local crowding which induces differentiation and subsequent delamination. Our work indicates that ID1-positive basal cells sustain progenitor states by coupling renewing proliferation with adhesion to the basement membrane (Figure 3 and 4). It is possible that *Id1* silenced progenitors are outcompeted due to additive defects in proliferation as well as adhesion. While our *in vitro* transcriptional profiling cannot temporally delineate activation of differentiation markers from down regulation of anchoring proteins, our *in vivo* characterization indicates that delamination proceeds differentiation, identifying suprabasal K10-positive cells retaining cell cycle or basal K5 progenitor marker expression (Figure 3). Interestingly, ID activity anchor neural stem cells to the extracellular matrix in the ventricular wall (Niola et al., 2012) suggesting that promotion of progenitor-to-niche adhesion is a common feature of ID-mediated transcriptional programs. Commitment to differentiation is a multistep process which requires the sequential activation of discrete transcriptional programs coordinating cell cycle exit and loss of progenitor cell adhesion with upregulation of differentiation markers. Here we unveil how fine-tuning of ID1 allows the developing epidermis to balance tissue expansion with epidermal stratification.

ID1 promotes proliferation of immortalized keratinocytes *in vitro* (Rotzer et al., 2006) but antagonizes HFSC activation *in vivo* (Genander et al., 2014). Here, we establish ID1 as a positive regulator of epidermal proliferation *in vivo* (Figure 4) and identify the interacting bHLH transcriptional factors TCF3 as a likely downstream effector. How ID1 promotes HFSC quiescence is unknown, but it is possible that HFSC-specific TCF heterodimerizing factors, lacking in the developing epidermis, directs the transcriptional response of TCF heterodimers in the absence of ID1. We were not able to identify additional heterodimerizing bHLH partners using cultured epidermal progenitors (Figure 5), suggesting that TCF3 could regulate epidermal transcriptional targets as a TCF-TCF homodimer. Homodimerization of TCF3 has been demonstrated to control cell fate in pluripotent stem cells as well as in the B-cell and myogenic lineage (Neuhold and Wold, 1993; Rao et al., 2020; Shen and Kadesch, 1995), and homodimers are more sensitive to ID1 inhibition compared to their TCF3-bHLH counterpart (Neuhold and Wold, 1993), increasing regulatory complexity in predicting the downstream transcriptional effects of bHLH activity. It would be interesting to functionally delineate the role of TCF dimerization in epidermal progenitors. Introduction of a C570A amino acid change in TCF3 is sufficient to inhibit homodimerization yet allowing TCF3-bHLH heterodimers (Benezra, 1994), and forced expression of TCF homodimers are reported (Neuhold and Wold, 1993; Rao et al., 2020; Sigvardsson et al., 1997) providing avenues for future systematic characterization of bHLH activity in the epidermis.

To help integrate the observed actions of ID1 in epidermal development, we incorporated transcriptional profiling of primary epidermal progenitor cells where *Id1* levels were altered. We identified a cohort of differentially expressed genes which positively or negatively correlated to *Id1* expression (Figure 5), many of which could be categorized as either mediators (*Col4a1, Col4a2, Cebpa*) or markers (*Dsg1a, Tgm3, Krt15*) of distinct aspects of epidermal differentiation. Transcription factor binding motif search revealed enrichment of bHLH binding sites in promoters of genes whose expression correlated to *Id1*, while uncorrelated genes were enriched for bZIP, Trp63, GRHL and T-box binding sites, all established mediators of distinct aspects of epidermal development (Lin et al., 2020; Lopez et al., 2009; Soares and Zhou, 2018).

Focusing on CEBPA, an interesting regulator of cell cycle exit and lineage commitment (Lopez et al., 2009; Nerlov, 2007), we confirmed enrichment of CEBPA in *shId1* compared to *shScr* targeted progenitors *in vivo* (Figure 5). Although functional bHLH binding E-boxes are described in the *Cebpa* enhancer (Cooper et al., 2015) and promoter (Pfurr et al., 2017; Soleimani et al., 2012; Yoon et al., 2015), we failed to activate or repress *Cebpa* reporter activity by manipulating TCF levels. We cannot exclude that ID1 act on *Cebpa* through TCF-independent mechanisms (Massari and Murre, 2000), or rule out additional unidentified cofactors required to activate *Cebpa* transcription in a TCF-dependent manner. It seems however more likely that the regulation of *Cebpa* is ID1-TCF independent and that CEBPA and TCF-mediated bHLH transcriptional programs act in parallel to cooperatively repress progenitor cell proliferation, thereby acting to safeguard progenitor fate transitions towards differentiation.

We noticed that overexpression of CEBPA reduced *Id1* mRNA and protein levels, suggesting upstream regulation of *Id1* by CEBPA in line with previous work demonstrating CEBPB-dependent regulation of ID proteins (Karaya et al., 2005; Mori et al., 2000; Saisanit and Sun, 1997). Exploiting a known BMP-sensing and pSMAD1/5-binding element in the *Id1* promoter (Genander et al., 2014; Korchynskyi and ten Dijke, 2002), we observed reduced BMP responsiveness upon differentiation or when CEBPA was introduced in progenitor cells and increased sensitivity in differentiated keratinocytes upon *Cebpa* silencing (Figure 7). BMP treatment is known to induce cell type specific transcriptional responses (Fessing et al., 2010) suggesting that expression of co-factors define context dependent SMAD-binding targets. CEBPA can interact with SMAD4 (Coyle-Rink et al., 2002; Zauberman et al., 2001), and redirect chromatin binding of SMAD1 during lineage specification (Trompouki et al., 2011). Our data suggest a model where BMP activity sustains progenitor states by inducing *Id1* while repressing *Cebpa*. Furthermore, we find that the presence or absence of CEBPA determines the ability of pSMAD1/5 to activate BMP-sensing chromatin, potentially diversifying the transcriptional output in response to BMP and thereby fine-tuning epidermal lineage progression. More specifically, our work indicates that CEBPA-positive basal progenitors are refractory to BMP-induced *Id1* promoter activation and consequently fated towards epidermal differentiation. We identify ID1 as a new transcriptional effector coordinating epidermal development.

## Material and Methods

### Mouse husbandry

Animals were housed in pathogen-free conditions according to the recommendations of the Federation of European Laboratory Animal Science Association. All animal experiments were approved by Stockholms djurförsöksetiska nämnd (ethical permis no N243/14, N116/16 and 14051-2019). Id1 floxed mice were previously described (Nam and Benezra, 2009), and Swiss mice were ordered (Janvier) for *in utero* lentiviral injections with shRNAs.

### Genotyping

Ear or tail biopsies were lysed overnight (o/n) at 55°C in DirectPCR lysis reagent (BioSite) in the presence of 0.1 mg/mL Proteinase K (Thermo Scientific). Lysis was stopped by heat inactivation (45min, 85°C). Taq Polymerase (5 U/*μ*L, final concentration: 0.05U/ *μ*L) was used for amplification in PCR buffer supplemented with 0.08 mM MgCl_2_, 0.2 mM dNTPs (all Invitrogen) and 0.4 *μ*M primers. PCR products were analyzed with 2% agarose gel.

### Antibody staining

Tissue samples used in this study were either fixed before paraffin embedding or snap frozen and fixed prior to antibody staining with 4% Formaldehyde (Sigma). Cells cultivated in chamber slides were subjected to the same fixative. For Immunohistochemistry antigen retrieval (R&D) was performed, then blocking (Bloxall and Vector stain kit) and finally antibody incubation as indicated by the manufacturer’s instructions. Samples used for immunofluorescence staining were cut at a thickness of 10 *μ*m, blocked with goat serum (2.5%) and BSA (1%) and permeabilized with 0.3% Triton X-100. When EdU was analyzed, the samples were treated with click chemistry prior to the antibody labelling (Click-iT, Thermo Fisher). Antibodies used for immunofluorescence staining are listed below. ID1 (Biocheck, #BCH-1, 1:500), Cleaved Caspase-3 (Cell Signaling Technology, #9661, 1:400), K10 (Biolegend, #905404, 1:1000), K5 (Biolegend, #905901, 1:1000), K14 (OriGene, #BP5009, 1:1000), Involucrin (Biolegend, #924401, 1:1000), TGM1 (Abcam, #ab103814, 1:1000), Flag (Sigma, F1804, 1:500), Cebpa (Cell Signaling Technology, #2295, 1:500). Antibodies used for immunoblotting are listed below: Gapdh (Abcam, #ab8245, 1:5000), Actin (Sigma, #A2228, 1:4000), Tubulin (Abcam, #ab7291, 1:5000). ID1 (Biocheck, #BCH-1, 1:1000), Flag (Sigma, F1804, 1:5000), Cebpa (Cell Signaling Technology, #2295, 1:1000) phospho-Smad1/5 (Cell Signaling Technology, #9516, 1:1000).

### RT-qPCR

Cells and tissue samples were collected in Trizol LS Reagent (Thermo Fisher). RNA was extracted using RNAeasy (Qiagen) or Direct-zol (Zymo), and treated with DNaseI (Qiagen). After the quality check, 100 ng of total RNA was used for cDNA synthesis using SuperScript IV Vilo (Thermo Fisher). RT-qPCR was run with selected primer pairs and SYBR Green on a ViiA 7 device or a 7500 fast system (both applied biosystems). HPRT was used as an internal control.

### RNAScope

RNA in situ hybridization to probe Tcf3, Tcf4, and Tcf12 was conducted using a Multiplex Fluorescent V2 kit from Advanced Cell Diagnostics (ACD), according to the manufacturer’s instruction. In brief, FFPE slides were baked at 60°C for 1hr, followed by deparaffinized with two washes of xylene (5 min for each) and two washes of 100% ethanol (2 min for each). The slides were then dried for 5min at 60°C, followed by incubation with hydrogen peroxide for 10 min at room temperature. The samples were then treated with protease at 40°C in the oven for 30 min, followed by incubation with Tcf3, Tcf4, and Tcf12 probes for 2 hrs at 40°C in the oven. After amplification with AMP 1, AMP 2, and AMP 3, each channel was developed by incubating HRP, fluorophore, and HRP blocker sequentially based on the channels of the probe that was incubated. The samples can next be counter-stained with DAPI (or antibodies if there’re other channels available). Images were taken with a Zeiss Axioplan microscope and analyzed with Zen blue software.

### Single-cell RNA-Sequencing

The datasets (E13_WT: 929 cells; E15_WT: 633 cells) provided by Fan et al. were obtained from the GEO database (GSE102086) transformed into a matrix and used as an input for the R package Seurat (version 2.3.0). For the E13 dataset, cells with less than 500 genes, more than 8500 genes, and more than 5% mitochondrial genes were excluded from further analyses. Seurat was used for clustering of the dataset. Count data was log-normalized with a scaling factor of 10,000 before identification of variable features. Variable features were identified utilizing variance stabilizing transformation and the top 2000 features were selected. Data scaling regressed out unwanted variation due to library size or mitochondrial gene content. Features were selected using PCA. Nearest neighbors were identified using the first 17 principal components and clusters identified using a resolution of 0,5. Utilizing the FindAllMarkers function keratinocytes were identified as being positive for *Krt5* and *Krt15* expression and negative for *Pdgfra, Vim, and Cd31* (154 cells at E13). Using the function subset re-analysis of the cells identified as keratinocytes was performed (variance stabilized transformed selecting top 2000 features, re-scaled data, re-ran PCA, performed nearest neighbor identification with 6 dimensions and clustering with a resolution parameter of 0,3) to visualize the expression of features of interest.

### In vitro establishment and expansion of primary epidermal progenitors

Primary epidermal progenitor cells were established and maintained according to published protocols (Nowak and Fuchs, 2009) using 3T3-J2 feeders and E-Low media. Differentiation was induced by increasing the calcium concentration to a final concentration of 1.5 mM.

### Production of high-titer lentivirus

Lentivirus was produced in 293TN cells (System Bioscience, LV900A-1) expanded in DMEM with 10% FBS as previously described (Beronja et al., 2010), using pMD2/pPAX packaging plasmids and calcium and the pLKO.1 back bone. Epidermal progenitors were infected by spinoculation (1100 x g for 30 minutes @ 37°C) in the presence of 40 *μ*g/mL hexadimethrine bromide (Beronja et al., 2010).

### In utero lentiviral injections

High titer lentivirus was injected into the amniotic cavity of E9.5 embryos in accordance with previous publications (Beronja et al., 2010) and with the help of INFINIGENE, Karolinska Institutet.

### Immunoprecipitation

The protein of interest (ID1, ID2, ID3, TCF3 or TCF4) was overexpressed by transfecting Flag-tagged constructs into mouse epidermal progenitor cells. Transfected cells were enriched using either puromycin or hygromycin, and the Flag-tag was precipitated using Sepharose beads pre-incubated with anti-FLAG or IgG antibody. Efficiency of precipitations was confirmed using western blot and if approved forwarded for mass spectrometry analyses to the Proteomics Biomedicum core facility, Karolinska Institutet. Interacting proteins were scored as hits based on the lack of signal in the IgG control, coverage and overall protein score.

### Luciferase Reporter Assay

CEBPa promoter (−2kb) or enhancer region (Cooper et al., 2015) was cloned into the pGL3 basic backbone (Promega). The construct containing part of the Id1 promoter region (336 bp) was previously described (Genander et al., 2014). Epidermal progenitor cells were cotransfected with the luciferase construct and Renilla-luciferin 2-monooxygenase RLuc (in a ratio of 1:10) using Lipofectamine LTX (Thermofisher). The empty pGL3 backbone was used for background control. Lysates were measured for luciferase activity and RLuc using the Dual Glo luciferase assay system on a Glomax reader (both Promega). To assess BMP sensitivity, cells were serum starved overnight and treated with BMP4 (200ng/mL) for 3 hours before lysed.

### RNA-Sequencing and Bioinformatic analyses

Total RNA was analyzed on a bioanalyzer and used for library preparation. Samples were sequences using a NovaSeq6000 platform (Illumina). A quality check (Fast QC/0.11.5) was run on raw sequencing reads. Raw sequencing reads were processed to obtain counts per genes for each sample. The EdgeR package was used to normalize for the RNA composition by finding a set of scaling factors for the library sizes that minimize the log-fold changes between the samples for most genes, using a trimmed mean of M values (TMM) between each pair of samples. Homer/4.10 was used to analyze promoters of genes and look for motifs that are enriched in the target gene promoters relative to all other promoters in the genome. Analyses, statistical computing and graphics were performed using R and in collaboration with NBIS (National Bioinformatics Infrastructure Sweden) at Karolinska Institutet.

### Statistical analyses

GraphPad Prism and R have been used for statistical analyses. Error bars show the SD or SEM as specified in the figure legends. Gene enrichment was analyzed using Gene Ontology (Ashburner et al., 2000).

## Acknowledgements

This study was supported by Vetenskapsrådet (2015-03215), Ragnar Söderbergs Stiftelse (M26/15), Stiftelsen för Strategisk Forskning (ICA14-0055), SSMFs Stora Anslag 2015 and Cancerfonden (2016/270). MG is a Ragnar Söderberg Fellow and Cancerfonden Junior Investigator. We are grateful for technical assistance from Karolinska Institutet core facilities INFINIGENE and Proteomics Biomedicum as well as NBIS (National Bioinformatics Infrastructure Sweden), SciLife, Stockholm. We thank members of the Genander lab for discussions.

## Author contributions

CGK, WY, DG, KVP and KHMM performed the experiments. CGK, WY, DG and MG analyzed the data. CGK, WY and MG designed the experiments and drafted the manuscript. All authors have proof-read and approved the final version of the manuscript.

