## Supplemental Information and Figures for "ID1 and CEBPA Coordinate Epidermal Progenitor Cell Differentiation"

**Supplemental Figures**

**Figure S1 ID1 is expressed in epidermal progenitor cells during skin development**

(A) Feature plot displaying cluster 1 and cluster 2 from E13 epidermal single-cell RNA-sequencing.

(B and C) Basal marker *Krt15* is enriched is cluster 1, whereas *Krtdap*, a marker of epidermal differentiation, is exclusively expressed in cluster 2.

(D) Gene Ontology analysis of biological processes using genes enriched in cluster 1 or 2.

(E) Protein classification of genes enriched in cluster 1 and cluster 2 respectively.

(F) Ridge plots of know markers of epidermal differentiation (*Mafb*, *Hes1* and *Klf4*) enriched in cluster 2.

(G) ID1 protein expression in developing epidermis at E15.5 and E16.5.

(H and I) Ridge and feature plots of *Id2* and *Id3* expression in E13 epidermis.

(J) Percentage of sequenced E13 epidermal progenitors expressing *Id1* alone, *Id1/2/3* together or are *Id1* negative.

(K) Number of differentially expressed genes found in *shId1* targeted cultured epidermal progenitors when compared to *shScr* cells.

Scale bars 75 µm (G E15.5) 50 µm (G E16.5).

**Figure S2 ID1 counteracts epidermal progenitor delamination**

(A) ID1 immunoreactivity is reduced in *Id1*$^{fl/fl}$ epidermis, but not *Id1*$^{+/fl}$, targeted with LV-CRE.

(B) *In vitro* transduction efficiency of *shScr* and *shId1* are comparable.

(C-E) Localization of transduced H2BGFP reporter positive cells at E14.5, E16.5 and E18.5 in *shScr* and *shId1* targeted epidermis.

(F and G) Distribution and quantification of LV-CRE targeted cells in *Id1*$^{+/fl}$ and *Id1*$^{fl/fl}$ at E14.5.

(H and I) Distribution and quantification of LV-CRE targeted K14-positive and K14-negative cells in $Id1^{+/fl}$ and $Id1^{fl/fl}$ at E16.5.

(J) Quantification of the number of cells/150um at E14.5 in *shScr* compared to *shId1* targeted epidermis.

(K and L) Immunoreactivity against cleaved caspase-3 (CC3) is not altered upon *Id1* silencing at E14.5.

(M) Measurement of epidermal thickness at E16.5 in *shScr* and *shId1*.

(N and O) Thickness of epidermis in $Id1^{+/fl}$ and $Id1^{fl/fl}$ targeted epidermis at E14.5 and E16.5.

(P) K10 spinous layer thickness in in $Id1^{+/fl}$ and $^{Id1fl/fl}$ targeted epidermis at E16.5.

Data are represented as mean ± SEM. $^{*}p < 0.05$ using unpaired t-test. Scale bars 50 µm.

**Figure S3 Progenitor cells devoid of ID1 co-express basal and differentiation markers**

(A) Immunoreactivity for K5 and K10 shows increased number of double-positive cells in $Id1^{fl/fl}$ skin targeted with LV-Cre compared to wild type epidermis.

(B) Reactome enrichment for genes differentially expressed at 24 hours of differentiation in *shId1* targeted cultured epidermal progenitors compared to *shScr* cells (>2xFC).

Scale bars 50 µm.

**Figure S4 Epidermal progenitor proliferation is positively regulated by ID1**

(A) Combined CRE and EdU immunoreactivity in $Id1^{+/fl}$ compared to $Id1^{fl/fl}$ epidermis at E16.5.

(B) *Id1* mRNA levels after Dox induction in cultured epidermal progenitors.

(C) ID1 protein is enriched in epidermal progenitors in doxycycline (Dox) treated cells compared to untreated cultures.

Scale bars 50 µm.

**Figure S5 Identification of ID1 gene signatures**

(A) Number of differentially expressed genes in ID1 overexpressing epidermal progenitor cells (0 hours) asked to differentiation (24 hours).

(B) Spinous markers expression is impaired at 24 hours of differentiation in ID1 overexpressing epidermal cells.

(C and D) Basal gene markers are not affected by ID1 overexpression.

(E and F) Overexpression of Flag-tagged ID1, ID2, ID3 TCF3 and TCF4 in cultured epidermal progenitors.

(G) HOMER motif analysis reveals enrichment of bZIP (basic leucine zipper domain), T-box, Trp63 and GRHL sequence motifs in promoters (+400bp) of Q2+Q4 genes when compared to all other expressed gene promoters.

(H) *Cebpa* mRNA levels increase with differentiation of epidermal progenitor cells.

(I) CEBPA protein levels are increased following *in vitro* differentiation of epidermal progenitors.

(J) Doxycycline dependent (3 days treatment) overexpression of CEBPA in epidermal progenitor cells.

(K and L) Cebpa promoter and enhancer luciferase reporter activity in s*hScr* and *shId1* targeted progenitors.

Data are represented as mean ± SEM. $^{***}p < 0.001$ using multiple unpaired t-test.

**Figure S6 TCF3/4/12 localize to the developing epidermis and regulate progenitor cell proliferation**

(A and B) *Tcf3*, *Tcf4* and *Tcf12* are uniformly expressed in both cluster 1 and 2 in E13 epidermis.

(C) Relative mRNA expression levels of *Tcf3*, *Tcf4* and *Tcf12* upon *in vitro* differentiation of epidermal progenitor cells.

(D) Knock down efficiency in epidermal progenitor cells *in vitro* using shRNAs targeting *Tcf3*, *Tcf4* or *Tcf12*.

(E) Expression profiles of differentiation markers in epidermal progenitor cells after silencing of *Id1*, *Tcf3*, *Tcf4* or *Tcf12*.

Data are represented as mean ± SD. $^{*}p < 0.05$ $^{**}p < 0.01$ $^{***}p < 0.001$ using multiple unpaired t-test.

**Figure S7 pSMAD1/5 activation of the *Id1* promoter is CEBPA dependent**

(A) mRNA induction of *Cebpa* upon doxycycline treatment.

(B) Relative silencing of *Cebpa* mRNA using two different shRNAs.

(C) *Id1* mRNA levels after silencing of *Cebpa* in epidermal progenitors.

(D) Protein levels of *Cebpa* after shRNA silencing in progenitor cells.

Data are represented as mean ± SD. $^{*}p < 0.05$ $^{**}p < 0.01$ $^{***}p < 0.001$ using multiple unpaired t-test.

**Suplemental Tables**

Table S1

Gene list of differentially expressed genes from Figure 5D, Q1 and Q3.

Table S2

Mass spectrometry data from overexpression of co-immunoprecipitation of 3x-Flag-ID1, ID2, ID3, TCF3 and TCF4. Proteins listed were identified in duplicate experiments, and not in IgG control samples.

Figure S1

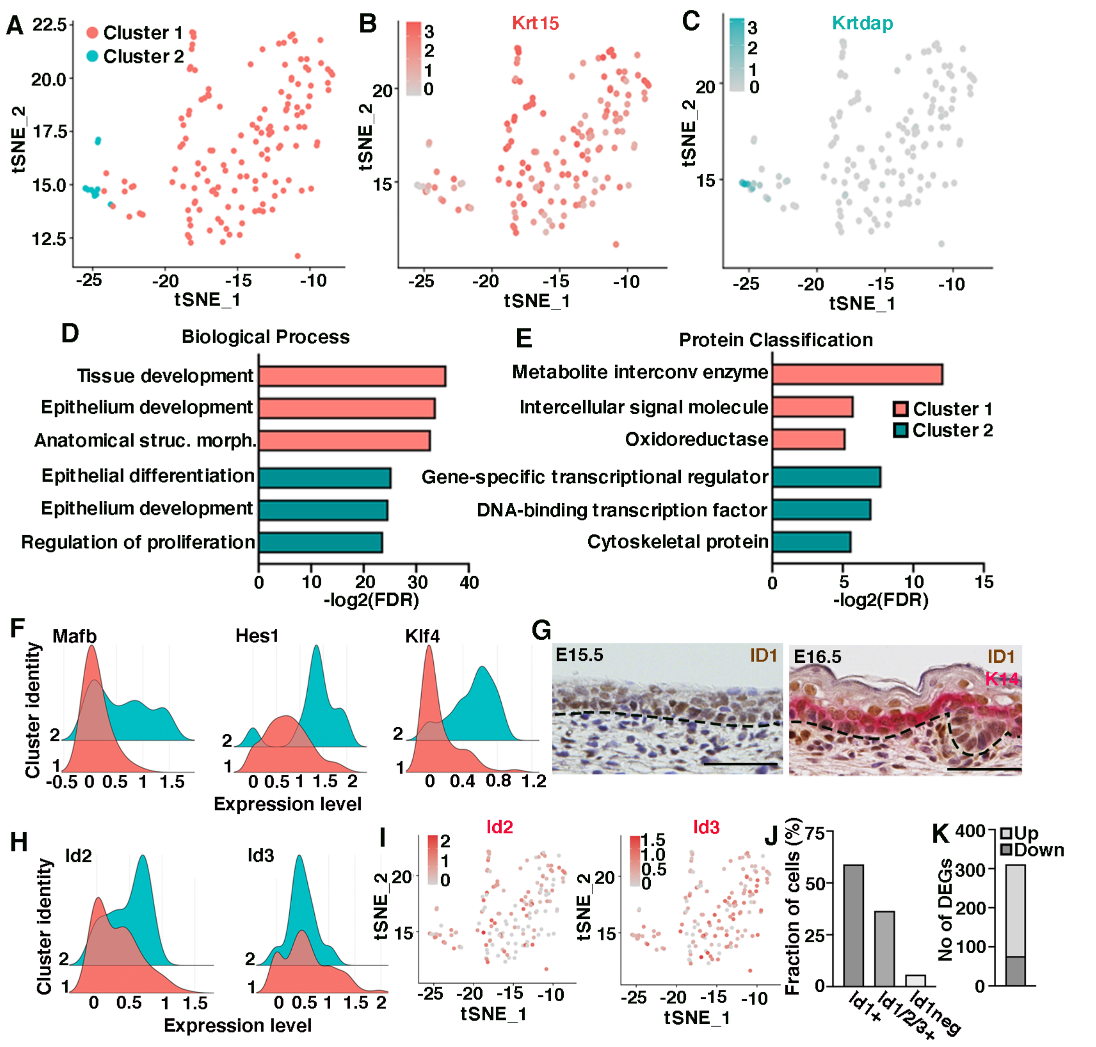
A
Cluster 1
Cluster 2
tSNE_2
tSNE_1
B
Krt15
C
Krtdap
D
Biological Process
Tissue development
Epithelium development
Anatomical struc. morph.
Epithelial differentiation
Epithelium development
Regulation of proliferation
-log2(FDR)
E
Protein Classification
Metabolite interconv enzyme
Intercellular signal molecule
Oxidoreductase
Gene-specific transcriptional regulator
DNA-binding transcription factor
Cytoskeletal protein
Cluster 1
Cluster 2
-log2(FDR)
F
Mafb
Hes1
Klf4
Cluster identity
Expression level
G
E15.5
ID1
E16.5
ID1
K14
H
Id2
Id3
Cluster identity
Expression level
I
Id2
Id3
tSNE_2
tSNE_1
J
Fraction of cells (%)
Id1+
Id1/2/3+
Id1neg
K
Up
Down
No of DEGs

Figure S2

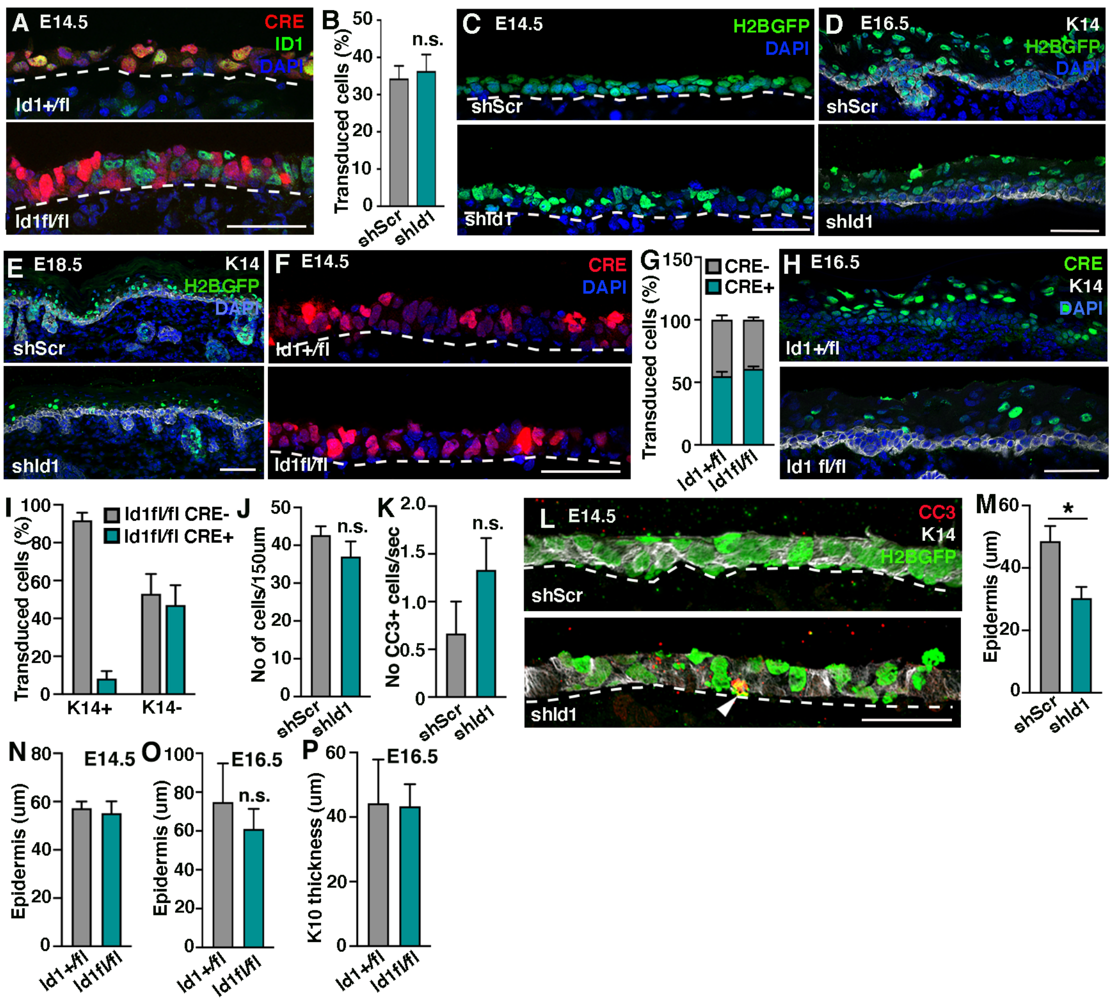

A E14.5
CRE
ID1
DAPI
Id1+/fl
Id1fl/fl
B
Transduced cells (%)
n.s.
shScr
shId1
C E14.5
H2BGFP
DAPI
shScr
shId1
D E16.5
K14
H2BGFP
DAPI
shScr
shId1
E E18.5
K14
H2BGFP
DAPI
shScr
shId1
F E14.5
CRE
DAPI
Id1+/fl
Id1fl/fl
G
CRE-
CRE+
Transduced cells (%)
Id1+/fl
Id1fl/fl
H E16.5
CRE
K14
DAPI
Id1+/fl
Id1 fl/fl
I
Id1fl/fl CRE-
Id1fl/fl CRE+
Transduced cells (%)
K14+
K14-
J
No of cells/150um
n.s.
shScr
shId1
K
No CC3+ cells/sec
n.s.
shScr
shId1
L E14.5
CC3
K14
H2BGFP
shScr
shId1
M
Epidermis (um)
*
shScr
shId1
N E14.5
Epidermis (um)
Id1+/fl
Id1fl/fl
O E16.5
Epidermis (um)
n.s.
Id1+/fl
Id1fl/fl
P E16.5
K10 thickness (um)
Id1+/fl
Id1fl/fl

**Figure S3**

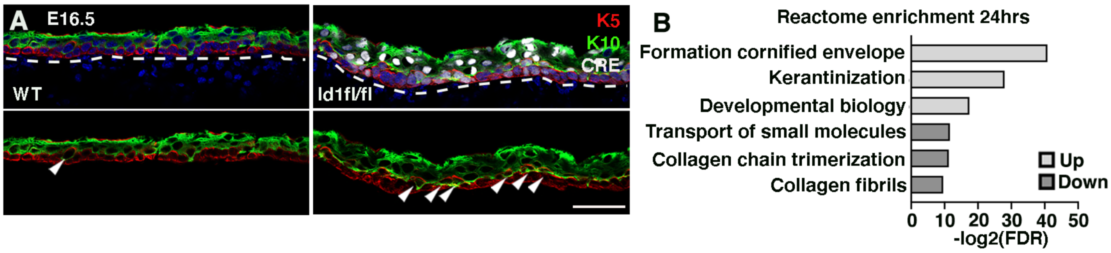
A
E16.5
WT
Id1fl/fl
K5
K10
CRE
B
Reactome enrichment 24hrs
Formation cornified envelope
Kerantinization
Developmental biology
Transport of small molecules
Collagen chain trimerization
Collagen fibrils
Up
Down
0 10 20 30 40 50
-log2(FDR)

### Figure S4

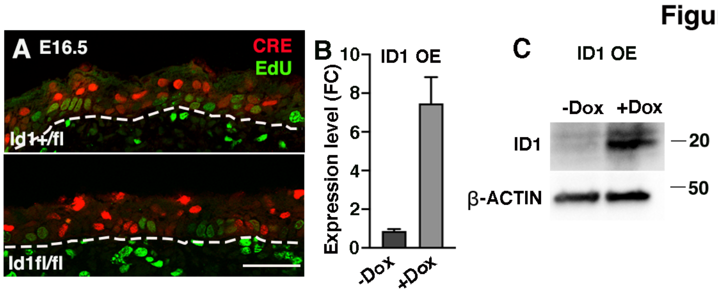
A E16.5
CRE
EdU
Id1+/fl
Id1fl/fl
B
ID1 OE
Expression level (FC)
10
8
6
4
2
0
-Dox
+Dox
C
ID1 OE
-Dox
+Dox
ID1
β-ACTIN
20
50

Figure S5

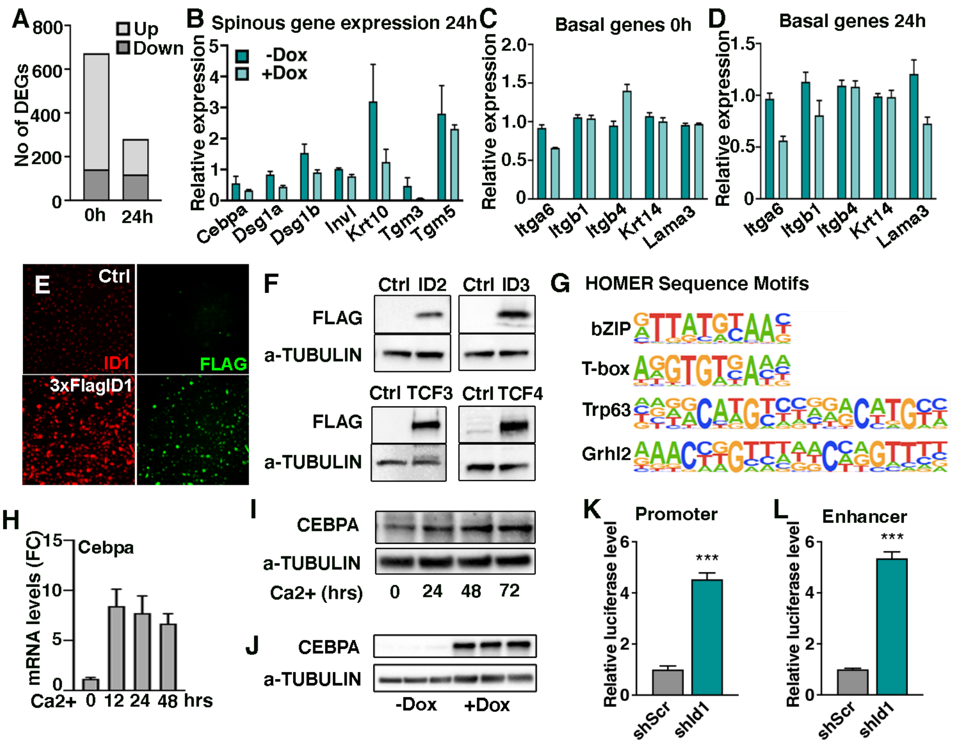
A
No of DEGs
800
600
400
200
0
Up
Down
0h
24h
B
Spinous gene expression 24h
Relative expression
-Dox
+Dox
Cebpa
Dsg1a
Dsg1b
Invl
Krt10
Tgm3
Tgm5
C
Basal genes 0h
Relative expression
Itga6
Itgb1
Itgb4
Krt14
Lama3
D
Basal genes 24h
Relative expression
E
Ctrl
ID1
FLAG
3xFlagID1
F
Ctrl ID2
Ctrl ID3
FLAG
a-TUBULIN
Ctrl TCF3
Ctrl TCF4
G
HOMER Sequence Motifs
bZIP
T-box
Trp63
Grhl2
H
mRNA levels (FC)
Cebpa
Ca2+ 0 12 24 48 hrs
I
CEBPA
a-TUBULIN
Ca2+ (hrs) 0 24 48 72
J
CEBPA
a-TUBULIN
-Dox
+Dox
K
Promoter
Relative luciferase level
***
shScr
shId1
L
Enhancer
Relative luciferase level
***
shScr
shId1

### Figure S6

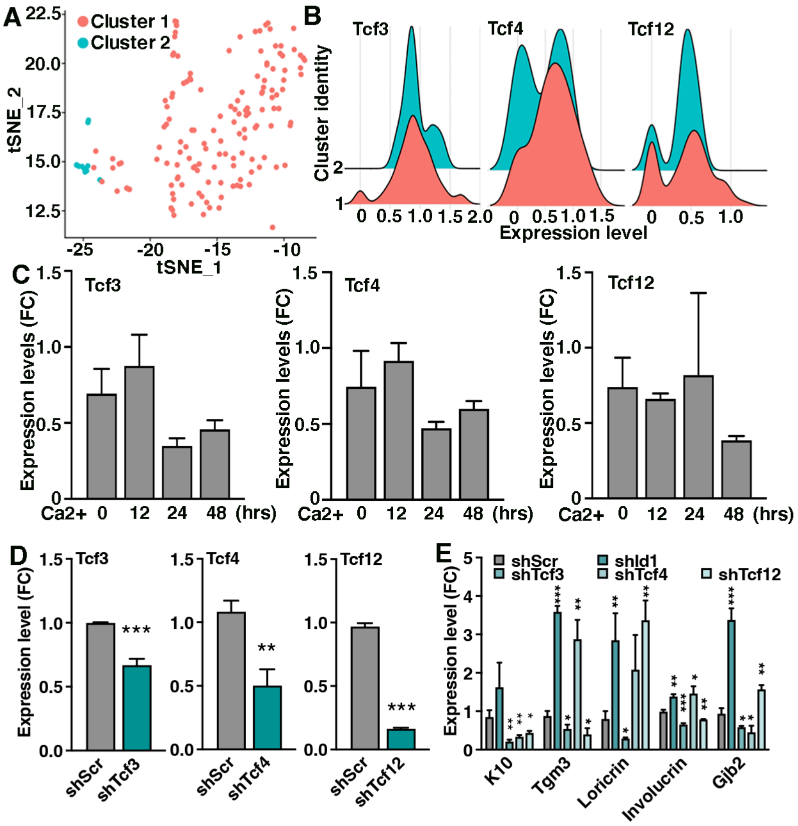

A
Cluster 1
Cluster 2
tSNE_2
tSNE_1
B
Tcf3
Tcf4
Tcf12
Cluster identity
Expression level
C
Tcf3
Tcf4
Tcf12
Expression levels (FC)
Ca2+ 0 12 24 48 (hrs)
D
Tcf3
Tcf4
Tcf12
Expression level (FC)
shScr
shTcf3
shTcf4
shTcf12
E
shScr
shId1
shTcf3
shTcf4
shTcf12
Expression level (FC)
K10
Tgm3
Loricrin
Involucrin
Gjb2

### Figure S7

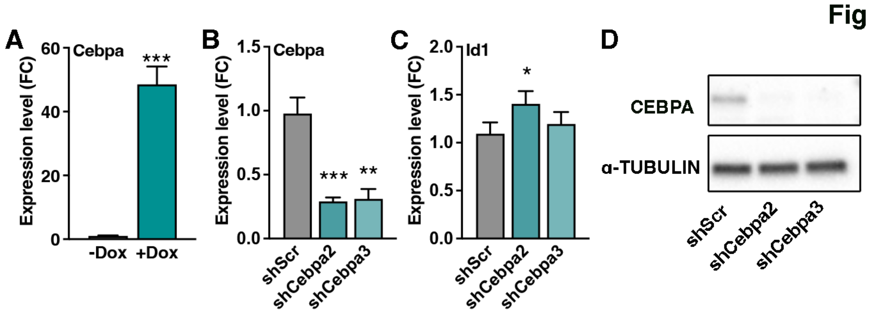
A
Cebpa
Expression level (FC)
-Dox
+Dox
B
Cebpa
shScr
shCebpa2
shCebpa3
C
Id1
D
CEBPA
α-TUBULIN
